# Sexual Dichromatism Drives Diversification Within a Major Radiation of African Amphibians

**DOI:** 10.1101/372250

**Authors:** Daniel M. Portik, Rayna C. Bell, David C. Blackburn, Aaron M. Bauer, Christopher D. Barratt, William R. Branch, Marius Burger, Alan Channing, Timothy J. Colston, Werner Conradie, J. Maximillian Dehling, Robert C. Drewes, Raffael Ernst, Eli Greenbaum, Václav Gvoždík, James Harvey, Annika Hillers, Mareike Hirschfeld, Gregory F.M. Jongsma, Jos Kielgast, Marcel T. Kouete, Lucinda P. Lawson, Adam D. Leaché, Simon P. Loader, Stefan Lötters, Arie van der Meijden, Michele Menegon, Susanne Müller, Zoltán T. Nagy, Caleb Ofori-Boateng, Annemarie Ohler, Theodore J. Papenfuss, Daniela Rößler, Ulrich Sinsch, Mark-Oliver Rödel, Michael Veith, Jens Vindum, Ange-Ghislain Zassi-Boulou, Jimmy A. McGuire

## Abstract

Theory predicts that sexually dimorphic traits under strong sexual selection, particularly those involved with intersexual signaling, can accelerate speciation and produce bursts of diversification. Sexual dichromatism (sexual dimorphism in color) is widely used as a proxy for sexual selection and is associated with rapid diversification in several animal groups, yet studies using phylogenetic comparative methods to explicitly test for an association between sexual dichromatism and diversification have produced conflicting results. Sexual dichromatism is rare in frogs, but it is both striking and prevalent in African reed frogs, a major component of the diverse frog radiation termed Afrobatrachia. In contrast to most other vertebrates, reed frogs display female-biased dichromatism in which females undergo color transformation, often resulting in more ornate coloration in females than in males. We produce a robust phylogeny of Afrobrachia to investigate the evolutionary origins of sexual dichromatism in this radiation and examine whether the presence of dichromatism is associated with increased rates of net diversification. We find that sexual dichromatism evolved once within hyperoliids and was followed by numerous independent reversals to monochromatism. We detect significant diversification rate heterogeneity in Afrobatrachia and find that sexually dichromatic lineages have double the average net diversification rate of monochromatic lineages. By conducting trait simulations on our empirical phylogeny, we demonstrate our inference of trait-dependent diversification is robust. Although sexual dichromatism in hyperoliid frogs is linked to their rapid diversification and supports macroevolutionary predictions of speciation by sexual selection, the function of dichromatism in reed frogs remains unclear. We propose that reed frogs are a compelling system for studying the roles of natural and sexual selection on the evolution of sexual dichromatism across both micro- and macroevolutionary timescales.

## INTRODUCTION

In *The Descent of Man and Selection in Relation to Sex*, Darwin (1871) observed that many closely related taxa differed primarily in secondary sexual characters and suggested that sexual selection plays a role in the diversification of species. The concept of speciation through sexual selection was later developed into a theory that links the coevolution of secondary sexual traits and mating preferences to premating reproductive isolation (Lande 1981, 1982; Kirkpatrick 1982; West-Eberhard 1983). This conceptual framework predicts that both the strength of sexual selection and the prevalence of sexually selected traits have a positive association with speciation rate (Lande 1981, 1982; West-Eberhard 1983; Barraclough et al. 1995). If divergence in secondary sexual characters and sexual selection are indeed major drivers of speciation, then clades exhibiting elaborate sexually dimorphic traits and/or elevated sexual selection should display higher species richness and increased diversification rates at macroevolutionary scales. Empirically, these predictions have mixed support across a range of sexually dimorphic traits that serve as proxies for sexual selection (reviewed in Kraaijeveld et al. 2011), indicating that macroevolutionary trends for traits involved with intersexual signaling and mate-choice may differ from those more strongly influenced by intrasexual or natural selection. For example, traits under ecological selection such as body size dimorphism have no consistent relationship with diversification (Kraaijeveld et al. 2011), whereas sexual dichromatism, a form of sexual dimorphism in which the sexes differ in color, often functions as a mate recognition signal and is associated with diversification in several taxonomic groups (Misof 2002; Stuart-Fox & Owens 2003; Alfaro et al. 2009; Kazancıo⍰lu et al. 2009; Wagner et al. 2012). Accordingly, sexual dichromatism has become a widely used proxy for studying the effects of sexual selection on speciation rate. However, phylogenetic comparative analyses explicitly testing for an association between sexual dichromatism and diversification have produced conflicting results, even in well-studied groups such as birds (Barraclough et al. 1995; Owens et al. 1999; Morrow et al. 2003; Phillimore et al. 2006; Seddon et al. 2013; Huang & Rabosky 2014) where sexual dichromatism plays an important role in signaling and mate-choice (Price 2008). Differences in methodologies and the spatial, temporal, and taxonomic scales among studies may partially explain these contrasting results. In particular, recent studies have highlighted concerns about the ability to distinguish between trait-dependent and trait-independent diversification scenarios using phylogenetic comparative methods (Rabosky & Goldberg 2015; Beaulieu & O’Meara 2016). In a broader sense, however, this disparity among studies may reflect more nuanced or novel mechanisms underlying the evolution of sexual dichromatism such that it does not consistently fit the conceptual framework of speciation by sexual selection. For instance, the striking sexual dichromatism in *Eclectus* parrots results from intrasexual competition to attract mates and intersexual differences in exposure to visual predators (Heinsohn et al. 2005). Likewise, the dynamic sexual dichromatism in frogs that form large breeding aggregations may serve to identify other competing males rather than to attract mates (Sztatecsny et al. 2012; Kindermann & Hero 2016; Bell et al. 2017b). Consequently, investigating the evolution of secondary sexual characters like dichromatism across ecologically diverse taxonomic groups is essential if we aim to generalize about the function of sexually dimorphic traits and better understand the roles of natural and sexual selection in biological diversification.

Secondary sexual traits are diverse and prevalent among anurans (frogs and toads), and include sexual size dimorphism, structures for acoustic signaling, nuptial pads, spines, elongated fingers, glands, and sexual dichromatism (Duellman & Trueb 1986). Sexual size dimorphism is common and occurs in >90% of species (Shine 1979; Han & Fu 2013), and body size evolution of the sexes is often attributed to fecundity for females and intrasexual competition, energetic constraints, agility, and predation for males (Salthe & Duellman 1973; Wells 1977; Shine 1979; Woolbright 1983; De Lisle & Rowe 2013; Han & Fu 2013; Nali et al. 2014). Male frogs of most species produce acoustic or vibrational signals to attract females, and these signals are essential for mate recognition and relaying social information (Ryan 1980; Gerhardt 1994; Gerhardt & Huber 2002). Structures like spines and tusks, which are present in males of many species, are used in male-male combat (e.g. McDiarmid 1975) whereas the diverse assortment of glands and nuptial pads, which are widespread in male frogs, are likely involved in courtship and amplexus (Duellman & Trueb 1986). In contrast to these widespread secondary sexual characters, anuran sexual dichromatism is rare, occurring in only ~2% of frog species (Bell & Zamudio 2012; Bell et al. 2017b). Behavioral studies in a handful of frog species indicate that these sexual color differences may be subject to natural selection as well as inter- and intrasexual selection (Mann & Cummings 2009; Sztatecsny et al. 2012). Sexual selection on coloration has historically been dismissed in frogs with the assumption that communication is predominately acoustic (reviewed in Starnberger et al. 2014); however, several studies document the importance of color signals in courtship behavior and mate-choice, even in nocturnal species (Gomez et al. 2009; Gomez et al. 2010; Jacobs et al. 2016; Yovanovich et al. 2017; Akopyan et al. 2018). If sexual dichromatism in frogs contributes to premating reproductive isolation, then sexually dichromatic clades may fit the conceptual framework of speciation by sexual selection and display higher species richness and increased diversification rates at macroevolutionary scales.

The Afrobatrachia frog radiation (Arthroleptidae, Brevicipitidae, Hemisotidae, Hyperoliidae) includes over 400 species distributed across sub-Saharan Africa that reflect much of the morphological, ecological and reproductive mode diversity present in anurans (Portik & Blackburn 2016). Afrobatrachian frogs display many unusual secondary sexual characters including the hair-like skin structures of male Hairy Frogs (*Trichobatrachus*), an elongate third digit in males (up to 40% of body length in *Arthroleptis* and *Cardioglossa*), extreme body size dimorphism (*Leptopelis, Chrysobatrachus, Breviceps*), and prominent pectoral glands (*Leptopelis*) or gular glands on the male vocal sac (Hyperoliidae). Afrobatrachian frogs also have the highest incidence of sexual dichromatism among anurans, which is striking and prevalent in many hyperoliid reed frogs (*Hyperolius, Heterixalus*). In sexually dichromatic hyperoliids, the difference in coloration is female-biased: both sexes exhibit a consistent juvenile coloration upon metamorphosis (termed Phase J), but at the onset of maturity sex steroids trigger a color and/or color pattern change in females (Phase F) whereas males typically retain the juvenile color pattern (Fig. 1) (Schiøtz 1967; Richards 1982; Hayes 1997; Hayes & Menendez 1999). In many dichromatic reed frog species adult males can also display the Phase F coloration, but this generally occurs in lower frequency than the Phase J morph (Fig. 1) (Schiøtz 1967, 1999; Amiet 2012; Kouamé et al. 2015; Portik et al. 2016a). The function of the ontogenetic color shift in female reed frogs is poorly understood (Bell & Zamudio 2012), but it may be similar to female-biased sexual dichromatism in other vertebrates, which can result from a reversal in mating system in which females compete for males (Andersson 1994) or sexual niche partitioning in which males and females use different habitats (Shine 1989; Heinsohn et al. 2005). Alternatively, distinct female color patterns in reed frogs may play a role in courtship and mate recognition at breeding sites where upwards of eight hyperoliid species congregate in a single night (Drewes & Vindum 1994; Kouamé et al. 2013; Portik et al. 2018). The link between sexual dichromatism and rapid speciation across disparate animal clades (Misof 2002; Stuart-Fox & Owens 2003; Alfaro et al. 2009; Kazancıo⍰lu et al. 2009; Wagner et al. 2012) demonstrates that when sexual dichromatism functions primarily as an intersexual signal under strong sexual selection, there are predictable outcomes on diversification rate. Therefore, as a first step towards understanding the potential function of dichromatism in hyperoliids, including the plausibility of intersexual signaling, we aim to assess whether this trait fits the predictions of speciation by sexual selection on a macroevolutionary scale.

**Figure 1.**
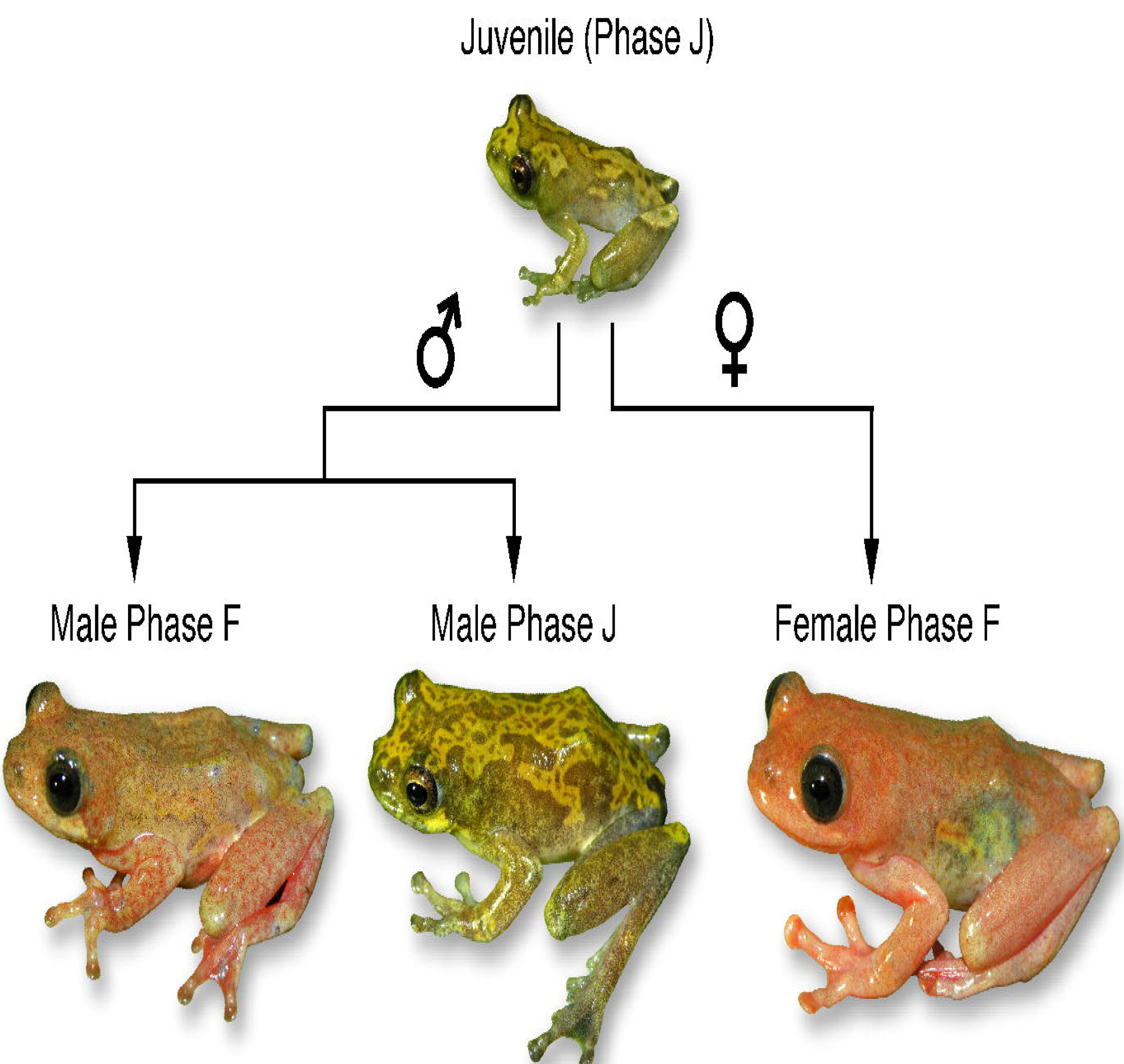
Illustration of ontogenetic color change occurring in hyperoliid frogs (*Hyperolius dintelmanni* shown) that underlies sexual dichromatism. In dichromatic species, females undergo a color change from Phase J to Phase F in response to steroid hormones at the onset of sexual maturity. Males retain the juvenile coloration (Phase J) or undergo a parallel change in color (to Phase F), however the proportion of the male color phases in populations varies across species. Secondary monochromatism can evolve from dichromatism through the loss of Phase J males (both sexes undergo an ontogenetic color change to Phase F) or through the loss of ontogenetic color change in both sexes (both sexes retain Phase J coloration at sexual maturity).

In this study, we reconstruct the evolutionary history of sexual dichromatism within Afrobatrachia and investigate whether diversification rate shifts in this continental radiation are associated with dichromatism. We produce a well-resolved species tree of Hyperoliidae from genomic data and greatly increased taxonomic sampling relative to previous studies (Wieczorek et al. 2000; Veith et al. 2009; Portik & Blackburn 2016). To explore the evolution of sexual dichromatism within the broader phylogenetic and biogeographic context of African frogs, we incorporate all published sequence data of Afrobatrachia to produce a robust, time-calibrated phylogeny. Using methods that improve the accuracy of character state reconstructions by accounting for trait and diversification rate heterogeneity (King & Lee 2015; Beaulieu & O’Meara 2016), we test the hypothesis that sexual dichromatism evolved repeatedly within Afrobatrachia (Veith et al. 2009). Finally, we estimate net diversification rates from our time-calibrated phylogeny using the hidden state speciation and extinction framework (HiSSE; Beaulieu & O’Meara 2016) and examine if diversification rate shifts occur within Afrobatrachia and if so, whether they are associated with the occurrence of sexual dichromatism. Given recent concerns raised about the ability to distinguish between trait-dependent and trait-independent diversification scenarios (Rabosky & Goldberg 2015; Beaulieu & O’Meara 2016), we assess the performance of available state-dependent diversification methods with a trait simulation study conducted using our empirical phylogeny.

## MATERIALS & METHODS

### Species Tree Estimation of Family Hyperoliidae

#### Taxonomic sampling

We included 254 hyperoliid samples in our sequence capture experiment with multiple representatives per species when possible. Although there are 230 currently recognized hyperoliid species (AmphibiaWeb, 2018), this family is in a state of taxonomic flux with recent studies recommending both the synonymy of species names and the splitting of species complexes (Rödel et al. 2002; Rödel et al. 2003; Wollenberg et al. 2007; Rödel et al. 2009; Rödel et al. 2010; Schick et al. 2010; Conradie et al. 2012; Dehling 2012; Channing et al. 2013; Conradie et al. 2013; Greenbaum et al. 2013; Liedtke et al. 2014; Loader et al. 2015; Portik et al. 2016a; Bell et al. 2017a; Conradie et al. 2018). We estimate that our sampling represents approximately 143 distinct hyperoliid lineages including 12 of 17 described hyperoliid genera. The five unsampled genera are either monotypic (*Arlequinus, Callixalus, Chrysobatrachus, Kassinula*) or species poor (*Alexteroon*, 3 spp.). Our sampling of the remaining genera is proportional to their recognized diversity and includes several known species complexes with lineages not reflected in the current taxonomy. We also sampled outgroup species from the following families: Arthroleptidae (7 spp), Brevicipitidae (1 sp), Hemisotidae (1 sp), and Microhylidae (1 sp). Museum and locality information for all specimens is provided in Table S1.

#### Sequence capture data and alignments

The full details of transcriptome sequencing, probe design, library preparation, sequence captures, and bioinformatics pipelines are described in Portik et al. (2016b), but here we outline major steps of the transcriptome-based exon captures. We sequenced, assembled, and filtered the transcriptomes of four divergent hyperoliid species and selected 1,260 orthologous transcripts for probe design. We chose transcripts 500–850 bp in length that ranged from 5–15% average pairwise divergence. Five additional nuclear loci (*POMC*, *RAG-1*, *TYR*, *FICD* and *KIAA2013*) were also incorporated based on published sequence data (Portik & Blackburn 2016). The final marker set for probe design included 1,265 genes from four species and 5060 individual sequences, with a total of 995,700 bp of target sequence. These sequences were used to design a MYbaits-3 custom bait library (MYcroarray, now Arbor Biosciences) consisting of 120 mer baits and a 2X tiling scheme (every 60 bp), which resulted in 60,179 unique baits. Transcriptomes, target sequences, and probe designs are available on Dryad (Portik et al. 2016c).

Genomic DNA was extracted using a high-salt extraction method (Aljanabi & Martinez 1997) and individual genomic libraries were prepared following Meyer & Kircher (2010) with modifications described in Portik et al. (2016b). Samples were pooled for capture reactions based on phylogenetic relatedness, and the combined postcapture libraries were sequenced on three lanes of an Illumina HiSeq2500 with 100 bp paired-end reads. Raw sequence data were cleaned following Singhal (2013) and Bi et al. (2012), and the cleaned reads of each sample were *de novo* assembled, filtered, and mapped as described in Portik et al. (2016b). The final filtered assemblies were aligned using MAFFT (Katoh et al. 2002, 2005; Katoh & Standley 2013) and trimmed using TRIMAL (Capella-Gutierrez et al. 2009). We enforced additional post-processing filters for alignments, including a minimum length of 90 bp and a maximum of 30% total missing data across an alignment, resulting in 1,047 exon alignments totaling 561,180 bp.

#### Species tree estimation

We used the sequence capture data set to estimate a species tree for Hyperoliidae using ASTRAL-III (Mirarab et al. 2014; Mirarab & Warnow 2015; Zhang et al. 2017), which uses unrooted gene trees to estimate the species tree. This method employs a quartet-based approach that is consistent under the multi-species coalescent process, and therefore appropriate for resolving gene tree discordance resulting from incomplete lineage sorting (Mirarab et al. 2014; Mirarab & Warnow 2015). This approach also allows for missing taxa in alignments, which were present in our sequence capture data and are problematic for other coalescent-based summary methods such as MP-EST (Liu et al. 2010). We kept samples with the most complete sequence data to collapse the alignments to a single representative per lineage and generated unrooted maximum likelihood (ML) gene trees with 200 bootstrap replicates for each locus using RAxML v8 (Stamatakis 2014) under the GTRCAT model. The set of 1,047 gene trees was used to infer a species tree with ASTRAL-III, and node support was assessed with 1) quartet support values, or local posterior probabilities computed from gene tree quartet frequencies, which also allows the calculation of branch lengths in coalescent units (Sayyari & Mirarab 2016), and 2) 200 replicates of multi-locus bootstrapping, where each of the 200 RAxML bootstrap trees per locus are used to infer a species tree and a greedy consensus tree is created from the 200 species trees to calculate percent support across nodes (Seo 2008).

### Evolutionary Relationships of Afrobatrachian Frogs

#### DNA barcoding

We obtained sequence data from the mitochondrial marker 16S ribosomal RNA (*16S*) for all samples included in the sequence capture experiment and for additional species we were unable to include in our sequence capture experiment due to insufficient DNA yield. Polymerase chain reactions (PCRs) were carried out in 12.5 μl volumes consisting of: 1.25 μl Roche 10x (500 mM Tris/HCl, 100 mM KCl, 50 mM (NH_4_)_2_ SO_4_, 20 mM MgCl_2_, pH =?8.3), 0.75 μl 25 mM MgCl_2_, 0.75 μl 2 mM dNTPs, 0.25 μl 10.0 μM forward primer, 0.25 μl 10.0 μM reverse primer, 8.40 μl H_2_O, 0.10 μl Taq, and 0.75 μl DNA. Amplification involved initial denaturation at 94°C for 4 min, followed by 35 cycles of 95°C for 60 s, 51°C for 60 s, 72°C for 90 s, and a final extension at 72°C for 7 min, using the primer pairs 16SA and 16SB (Palumbi et al. 1991). The PCR amplifications were visualized on an agarose gel, cleaned using ExoSAP-IT, and sequenced using BigDye v3.1 on an ABI3730 sequencer (Applied Biosystems). Newly generated sequences are deposited in GenBank (Accession numbers: XXXXX).

#### GenBank data

To expand our taxonomic sampling we included all available published sequence data of Afrobatrachian frogs. We built a molecular data matrix using the five nuclear loci included in our captures (*FICD*, *KIAA2013*, *POMC*, *TYR*, and *RAG-1*) and the mtDNA marker *16S*. This resulted in the inclusion of 30 additional hyperoliids and 130 arthroleptid, brevicipitid, and hemisotid species, though many are represented solely by *16S* mtDNA data. Nuclear loci were aligned using MUSCLE (Edgar 2004), and *16S* sequences were aligned using MAFFT with the E-INS-i strategy (Katoh et al. 2002; Katoh et al. 2005). The final concatenated alignment of the expanded taxonomic data set consisted of 283 taxa and 3,991 bp, with 36% total missing data.

#### Phylogenetic methods and divergence dating analyses

We reconstructed the phylogenetic relationships of Afrobatrachian frogs from the five nuclear gene and mtDNA data set using a maximum likelihood (ML) approach in RAXML v8 (Stamatakis 2014). To preserve the relationships inferred with our species tree analyses of sequence capture loci, the sequence capture hyperoliid species tree containing 153 taxa (143 ingroup and 10 outgroup taxa) was used as a partial constraint tree for the ML analysis of the 283-taxon alignment of six loci. A preliminary step for our divergence dating analyses in BEAST v1.8.1 (Drummond et al. 2012) involved transforming the ML Afrobatrachian frog tree to an ultrametric tree, and for this we used penalized likelihood with the ‘chronopl’ function of APE (Sanderson 2002; Paradis et al. 2004), setting age bounds to allow divergence times to be compatible with our BEAST calibration priors. We performed analyses using a fixed ultrametric starting tree topology by removing relevant operators acting on the tree model. We used four secondary calibration points with normal distributions to constrain the most recent common ancestors (MRCAs) of Afrobatrachia to 80 Ma ± 5 SD, (Hemisotidae + Brevicipitidae) to 50 Ma ± 5 SD, Arthroleptidae to 40 Ma ± 5 SD, and Hyperoliidae to 40 Ma ± 5 SD, based on published age estimates of Afrobatrachian frogs (Roelants et al. 2007; Kurabayashi & Masayuki 2013; Loader et al. 2014; Portik & Blackburn 2016). We used the Yule model of speciation as the tree prior, applied an uncorrelated relaxed lognormal clock, and ran two analyses for 30,000,000 generations sampling every 3,000 generations. Runs were assessed using TRACER v1.5.0 (Rambaut et al. 2013) to examine convergence, and a maximum clade credibility tree with median heights was created from 7,500 trees after discarding a burn-in of 2,500 trees.

### Evolution of Sexual Dichromatism and State-Dependent Diversification

We scored the presence or absence of sexual dichromatism for hyperoliid species in our data set from multiple sources, including publications (Schiøtz 1967, 1999; Channing 2001; Channing & Howell 2006; Wollenberg et al. 2007; Rödel et al. 2009; Veith et al. 2009; Amiet 2012; Bell & Zamudio 2012; Channing et al. 2013; Conradie et al. 2013; Portik et al. 2016a), examination of museum specimens (Portik 2015), and the collective field observations from all authors. A species was considered sexually dichromatic if adult females and adult males exhibit distinct color patterns (Phase F and Phase J, respectively), which also includes species with variation in male color phase (males display Phase J and Phase F). A summary of the sexual dichromatism data is provided in Table S2.

We reconstructed the evolution of sexual dichromatism on the time-calibrated phylogeny of Afrobatrachia using Bayesian ancestral state reconstruction in BEAST (Drummond et al. 2012), and also with hidden state speciation and extinction analyses using the R package HISSE (Beaulieu & O’Meara 2016) (described below). We performed Bayesian ancestral state reconstructions using several combinations of clock and character models in BEAST v1.8.1 (Drummond et al. 2012) following King and Lee (2015). The topology and branch lengths were fixed by removing all tree operators, and sexual dichromatism was treated as a binary alignment. Because all outgroup families are monochromatic, we fixed the root state by adding a placeholder monochromatic taxon to the root with a zero branch length, creating a hard prior distribution on the root where P(monochromatic root) = 1, and P(dichromatic root) = 0. A stochastic Mk model of character evolution (Lewis 2001) was used with symmetrical (Mk1) or asymmetrical (Mk2) transition rates between states, and each character model was analyzed using a strict clock model (enforcing a homogenous trait rate) and a random local clock model (allowing for heterotachy), resulting in four analysis combinations. The analyses involving the random local clock allowed estimation of the number and magnitude of rate changes using MCMC. All analyses were run for 200 million generations with sampling every 20,000 generations, resulting in 10,000 retained samples. We used the marginal likelihood estimator with stepping stone sampling, with a chain length of 1 million and 24 path steps, to estimate the log marginal likelihood of each run (Baele et al. 2012, 2013). We performed five replicates per analysis to ensure consistency in the estimated log marginal likelihood, and subsequently compared the four different analyses using log Bayes factors, calculated as the difference in log marginal likelihoods, to select the best fit clock model and character model combination. We summarized transitions between character states and created consensus trees to estimate the posterior probabilities of a character states across nodes.

We performed hidden state speciation and extinction (HiSSE) analyses using the R package HISSE (Beaulieu & O’Meara 2016) to identify if sexual dichromatism in Afrobatrachian frogs is associated with increased diversification rates, and to reconstruct ancestral states while accounting for transition rate and diversification rate heterogeneity. The HiSSE model builds upon the popular binary-state speciation and extinction (BiSSE) model (Maddison et al. 2007) by incorporating ‘hidden states’ representing unmeasured traits that could impact the diversification rates estimated for states of the observed trait. The HiSSE model is therefore able to account for diversification rate heterogeneity that is not linked to the observed trait, while still identifying trait-dependent processes. The HiSSE framework also includes a set of null models that explicitly assume the diversification process is independent from the observed trait, without constraining diversification rates to be homogenous across the tree. The inclusion of these character-independent models circumvents a significant problem identified in the BiSSE framework, in which the simple ‘null’ model of constant diversification rates is typically rejected in favor of trait-dependent diversification when diversification rate shifts unrelated to the trait occur in the phylogeny (Rabosky & Goldberg 2015; Beaulieu & O’Meara 2016). The improved character-independent diversification models, referred to as CID-2 and CID-4, contain the same number of diversification rate parameters as the BiSSE and HiSSE models, respectively. We fit 26 different models to our sexual dichromatism data set (Table 1): six represent BiSSE-like models, four are variations of the CID-2 model, five are variations of the CID-4 model, nine are various HiSSE models with two hidden states, and two are HiSSE models with a single hidden state. Within each of these classes, the models vary mainly in the number of distinct transition rates (q), extinction fraction rates (ε), and net turnover rates (τ), and the most complex HiSSE model includes four net turnover rates, four extinction fraction rates, and eight distinct transition rates. We enforced a monochromatic root state for all models and evaluated the fit of the 26 models using AIC scores, ΔAIC scores, and Akaike weights (ω_i_) (Burnham & Anderson 2002). From the best-fit model, we estimated confidence intervals for relevant parameters and transformed ε and τ to obtain speciation (λ), extinction (μ), and net diversification rates using the ‘SupportRegion’ function in HISSE (Beaulieu & O’Meara 2016). We performed ancestral state estimations for each of the 26 models using the marginal reconstruction algorithm implemented in the ‘MarginRecon’ function of HISSE, again enforcing a monochromatic root state. Our final estimation and visualization of diversification rates and node character states on the Afrobatrachian phylogeny took model uncertainty into account by using the model averaging approach described by Beaulieu and O’Meara (2016), such that model contributions to rates and states were proportional to their likelihoods.

**Table 1.** Summary of models and fits using hidden state speciation and extinction analyses, with sexual dichromatism as the observed state. The top ranked model is highlighted in bold.

| Description | Model | loglik | AIC | ΔAIC | wtAIC | Hidden States | State-dependent | Net turnover rates | Extinction fraction rates | Distinct transition rates |
| --- | --- | --- | --- | --- | --- | --- | --- | --- | --- | --- |
| **HiSSE** | **19** | **-1033.5** | **2082.9** | **0.0** | **0.445** | **yes** | **yes** | **4** | **1** | **3** |
| HiSSE | 15 | -1026.0 | 2084.0 | 1.0 | 0.266 | yes | yes | 4 | 4 | 8 |
| HiSSE | 18 | -1029.7 | 2085.4 | 2.5 | 0.130 | yes | yes | 4 | 1 | 8 |
| CID-4 | 14 | -1035.3 | 2086.5 | 3.6 | 0.075 | yes | no | 4 | 1 | 3 |
| CID-2 | 8 | -1031.7 | 2087.3 | 4.4 | 0.049 | yes | no | 2 | 2 | 8 |
| HiSSE | 16 | -1033.5 | 2088.9 | 6.0 | 0.022 | yes | yes | 4 | 4 | 3 |
| HiSSE | 25 | -1036.1 | 2092.2 | 9.3 | 0.004 | only for state 0 | yes | 3 | 3 | 4 |
| CID-4 | 12 | -1035.3 | 2092.5 | 9.6 | 0.004 | yes | no | 4 | 4 | 3 |
| CID-4 | 26 | -1029.7 | 2093.4 | 10.4 | 0.002 | yes | no | 4 | 4 | 9 |
| CID-2 | 10 | -1036.2 | 2094.4 | 11.5 | <0.001 | yes | no | 2 | 1 | 8 |
| HiSSE | 24 | -1041.1 | 2102.3 | 19.3 | <0.001 | only for state 1 | yes | 3 | 3 | 4 |
| BiSSE-like | 2 | -1046.3 | 2102.6 | 19.7 | <0.001 | no | yes | 2 | 1 | 2 |
| BiSSE-like | 1 | -1046.3 | 2104.6 | 21.7 | <0.001 | no | yes | 2 | 2 | 2 |
| BiSSE-like | 3 | -1053.5 | 2115.1 | 32.1 | <0.001 | no | no | 1 | 1 | 2 |
| HiSSE | 22 | -1055.7 | 2121.5 | 38.5 | <0.001 | yes | yes | 4 | 1 | 3 |
| HiSSE | 21 | -1053.6 | 2127.2 | 44.3 | <0.001 | yes | yes | 4 | 1 | 8 |
| CID-4 | 13 | -1060.2 | 2132.4 | 49.4 | <0.001 | yes | no | 4 | 1 | 1 |
| HiSSE | 20 | -1061.9 | 2135.7 | 52.8 | <0.001 | yes | yes | 4 | 1 | 1 |
| HiSSE | 17 | -1060.4 | 2138.7 | 55.8 | <0.001 | yes | yes | 4 | 4 | 1 |
| CID-4 | 11 | -1060.4 | 2138.8 | 55.8 | <0.001 | yes | no | 4 | 4 | 1 |
| CID-2 | 9 | -1068.1 | 2144.2 | 61.3 | <0.001 | yes | no | 2 | 1 | 1 |
| CID-2 | 7 | -1068.1 | 2146.2 | 63.3 | <0.001 | yes | no | 2 | 2 | 1 |
| BiSSE-like | 4 | -1071.4 | 2152.8 | 69.8 | <0.001 | no | yes | 2 | 2 | 1 |
| HiSSE | 23 | -1073.9 | 2153.7 | 70.8 | <0.001 | yes | yes | 4 | 1 | 1 |
| BiSSE-like | 5 | -1075.2 | 2158.3 | 75.4 | <0.001 | no | yes | 2 | 1 | 1 |
| BiSSE-like | 6 | -1077.6 | 2161.2 | 78.3 | <0.001 | no | no | 1 | 1 | 1 |

#### Simulated traits and state-dependent diversification

The identified bias towards the detection of trait-dependent diversification in the BiSSE framework (Rabosky & Goldberg 2015; Beaulieu & O’Meara 2016) prompted us to investigate if a similar outcome would be detected in our data set, and whether the HiSSE framework could improve our ability to distinguish whether the observed states are correlated with diversification rates. One concern raised by Beaulieu and O’Meara (2016) is that neutrally evolving traits simulated on trees generated from a complex heterogenous rate branching process can lead to false signals of trait-dependent diversification, signifying the HiSSE framework may be sensitive to particular types of tree shapes.

To evaluate the performance of these methods given our empirical tree topology, we simulated neutrally evolving traits on the Afrobatrachian frog phylogeny and tested for trait-dependent diversification using both the BiSSE and HiSSE frameworks. We conducted independent simulations of a binary trait with the ‘sim.history’ function in R package PHYTOOLS (Revell 2012), using the unequal rates *q*-matrices obtained from our empirical data, enforcing a root state of trait absence. We required a minimum of 10% of taxa to exhibit the derived state and conducted simulations until we obtained 1,500 replicates meeting this criterion. We used maximum likelihood to fit a BiSSE model and the typical ‘null’ model with equal speciation and extinction rates to each simulated trait using the R package DIVERSITREE (Maddison et al. 2007; FitzJohn et al. 2009; FitzJohn 2012). We performed likelihood ratio tests and calculated ΔAIC scores to determine if the constraint model could be rejected with confidence (ΔAIC > 2 or p-value < 0.05), and summarized the number of instances each of the two models was favored across the simulations. We analyzed the simulated data in the HiSSE framework as with our empirical data, but with a reduced set of five models representing each major category of model. This reduced model set included two BiSSE-like models that differed only in the constraint of τ, a CID-2 and CID-4 model, and a HiSSE model with two hidden states, three transition rates, equal τ, and distinct τ. We set a probability of one for trait absence at the root state to match the manner in which traits were simulated and evaluated the fit of the 5 models using ΔAIC scores and Akaike weights (ω_i_) for each simulation, using a threshold of ΔAIC greater than two to favor a model. Specifically, we were interested in whether the unconstrained BiSSE-like or HiSSE models were favored, resulting in the detection of a false pattern of trait-dependent diversification, or if the CID-2 or CID-4 models were selected, capturing the expected pattern where the diversification process was independent from trait evolution.

## RESULTS

### Phylogenetic Relationships

The sequence capture data set consisting of 1,047 loci and 561,180 bp produced a well-resolved species tree with a normalized quartet score of 0.877 and only eight of 150 nodes (5%) with quartet scores below 0.9 (Fig. S1). The multilocus bootstrapping analysis produced similar results with low support for only seven nodes, six of which also received low support in the species tree analyses (Fig. S1). The higher-level relationships recovered in the species tree are largely congruent with those recovered by Portik and Blackburn (2016), though we found strong support for a different placement of the genus *Acanthixalus* as sister to the clade containing *Semnodactylus, Paracassina, Phlyctimantis*, and *Kassina*, which together are sister to all other hyperoliids. Our improved sampling provides the first comprehensive assessment of evolutionary relationships within the speciose genera *Afrixalus* and *Hyperolius*. The monophyly of *Hyperolius* is supported, however *Afrixalus* is paraphyletic, and we found a sister relationship between the Ethiopian-endemic *A. enseticola* and the Malagasy-Seychelles species of *Heterixalus* and *Tachycnemis*, which are in turn sister to all remaining *Afrixalus* (Fig. S1). We found strong support for the southern African species *Hyperolius semidiscus* as sister to all other lineages in the genus, and furthermore recovered *H. parkeri*, *H. lupiroensis* and the *H. nasutus* complex as sister to two larger subclades of *Hyperolius* (Clades 1 and 2; Figs. 2, S1). In an effort to distinguish the main division within Hyperoliidae, we recognize the subfamilies Kassininae Laurent, 1972 and Hyperoliinae Laurent, 1943 and define the content within each on the basis of our species tree analysis as follows: 1) Kassininae: *Acanthixalus, Kassina, Paracassina, Phlyctimantis*, and *Semnodactylus*, 2) Hyperoliinae: *Afrixalus*, *Cryptothylax*, *Heterixalus*, *Hyperolius*, *Morerella*, and *Opisthothylax* (Fig. 2). We retain previous subfamily assignments for genera not sampled in our molecular study (Hyperoliinae: *Alexteroon*, *Arlequinus*, *Callixalus*, *Chrysobatrachus*, *Kassinula*), which should be included in future phylogenetic studies to confirm these placements.

**Figure 2.**
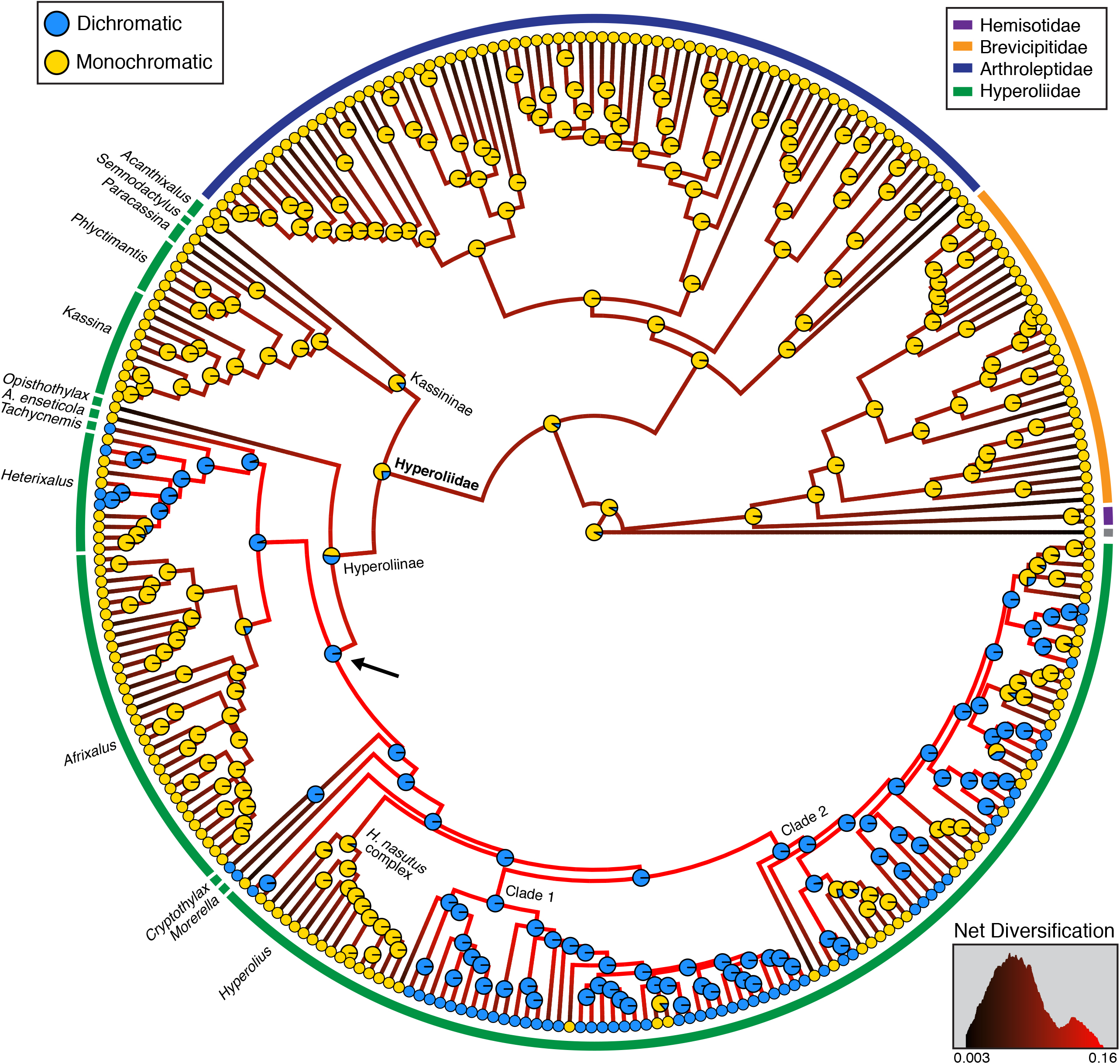
Ancestral state reconstruction of sexual dichromatism in Afrobatrachian frogs from HiSSE using model averaging to account for uncertainty in both models and reconstructions. Circles at tips and nodes are colored by state (yellow: monochromatic, blue: sexually dichromatic), with node pie charts indicating probability of a state assignment. Branches are colored using a gradient from the model-averaged net diversification rate, with black representing the slowest rate and red representing the fastest rate. The arrow highlights the node identified as displaying an unambiguous transition from monochromatism to sexual dichromatism.

The phylogenetic analyses of the Afrobatrachia supermatrix produced family-level relationships consistent with previous analyses (Pyron & Wiens 2011; Portik & Blackburn 2016; Feng et al. 2017), including a sister relationship between Hyperoliidae and Arthroleptidae, and between Hemisotidae and Brevicipitidae (Fig. S2). This expanded taxonomic data set also included improved sampling for the Malagasy hyperoliid genus *Heterixalus* and the *Hyperolius nasutus* complex, for which we recovered results consistent with Wollenberg et al. (2007) and Channing et al. (2013), respectively. We recovered an Eocene age for the time to most recent common ancestor (TMRCA) of the families Hyperoliidae, Arthroleptidae and Brevicipitidae, of approximately 42.4 Ma (95% HPD: 39.1–51.6 Ma), 45.4 Ma (95% HPD: 35.5–48.7 Ma), and 41.8 Ma (95% HPD: 33.9–49.8 Ma) (Fig. S2). These dates are younger than previous estimates (Roelants et al. 2007; Loader et al. 2014) but are consistent with more recent multilocus analyses (Portik & Blackburn 2016; Feng et al. 2017). We found diversification events began within Hyperoliinae approximately 35.8 Ma (95% HPD: 30.1–41.6 Ma) and within Kassininae approximately 38.3 Ma (95% HPD: 30.9–45.6 Ma). The majority of speciation events within *Hyperolius* occurred from the late Miocene to the Plio-Pleistocene (Fig. S2).

### Evolution of Sexual Dichromatism and State-Dependent Diversification

We found sexual dichromatism occurs in 60 of the 173 (34%) hyperoliid frog species included in our analyses. The Bayesian ancestral state reconstructions using four model combinations produced different patterns of character reconstructions and rates of trait evolution (Fig. S3). Overall, we found models with asymmetric character transition rates (Mk2) outperformed those with symmetric rates (Mk1), regardless of the clock model used (log-transformed Bayes factor range: 3.40–10.39; Table 2). The analysis with the highest log-marginal likelihood incorporated the relaxed local clock (RLC) Mk2 model, though it was not a significantly better fit than the simpler strict clock (SC) Mk2 model (log-transformed Bayes factor of 0.10) indicating low variation in lineage-specific evolutionary rates for this trait. The ancestral character reconstructions of the SC Mk2 indicated a median of two transitions from monochromatism to dichromatism. However, the character reconstructions revealed that this estimate is due to low posterior probability estimates for two critical nodes within Hyperoliinae (Fig. S3), and the presence of dichromatism at either one or both of these nodes would result in a single origin of sexual dichromatism. By contrast, the SC Mk2 model demonstrated approximately 27 independent losses of sexual dichromatism within hyperoliids, including losses in entire groups (*Afrixalus*, *H. nasutus* complex) and numerous reversals within *Heterixalus* and *Hyperolius* (notably in Clade 2) (Fig. S4).

**Table 2.** Summary of trait rate analyses for the evolution of sexual dichromatism. For transition summaries, monochromatism is coded as ‘0’ and dichromatism as ‘1’. The average number of rate shifts is provided, rather than median.

| Trait Rate Analysis | Log<br>Marginal<br>Likelihood | Transitions<br>0 > 1 | Transitions<br>1 > 0 | Rate<br>Shifts | Rate<br>Minimum | Rate<br>Maximum |
| --- | --- | --- | --- | --- | --- | --- |
| Strict Clock Mk1 | -92.12 | 15 | 7 | – | – | – |
| Strict Clock Mk2 | -81.83 | 2 | 27 | – | – | – |
| Relaxed Local Clock Mk1 | -85.10 | 16 | 8 | 3.3 | 0.001 | 0.019 |
| Relaxed Local Clock Mk2 | -81.73 | 2 | 27 | 0.8 | 0.013 | 0.016 |

Our hidden state speciation and extinction analyses revealed that a HiSSE model with four net turnover rates, equal extinction fraction rates, and three distinct transition rates was the best-fit model (Model 19, Table 1). The second and third ranked models (ΔAIC of 1.0, 2.5) were also HiSSE models that varied in the number of extinction fraction rates or the number of transition rates. Together these three HiSSE models accounted for 84.1% of the model weight (Table 1) and support a signal of state-dependent diversification in which sexual dichromatism and a hidden state are associated with diversification rates. The net diversification rates inferred using the best-fit model were nearly twice as high in sexually dichromatic lineages (0.157) as compared to monochromatic lineages (0.091) in the absence of the hidden state (Fig. 3), and the combination of the hidden state and dichromatism or monochromatism resulted in much lower net diversification rate estimates (0.02 and <0.001, respectively). The trait reconstructions for all four state combinations indicated that the dichromatism plus hidden state combination was only present in the MRCA of two genera (*Cryptothylax*, *Morerella*) (Fig. S5), and was associated with a markedly lower diversification rate (Fig. 3). The model-averaged ancestral state reconstructions demonstrated strong support for a single origin of sexual dichromatism and 25 independent reversals to monochromatism within Hyperoliinae, with reversal patterns similar to the Mk2 analyses (Figs. 2, S5).

**Figure 3.**
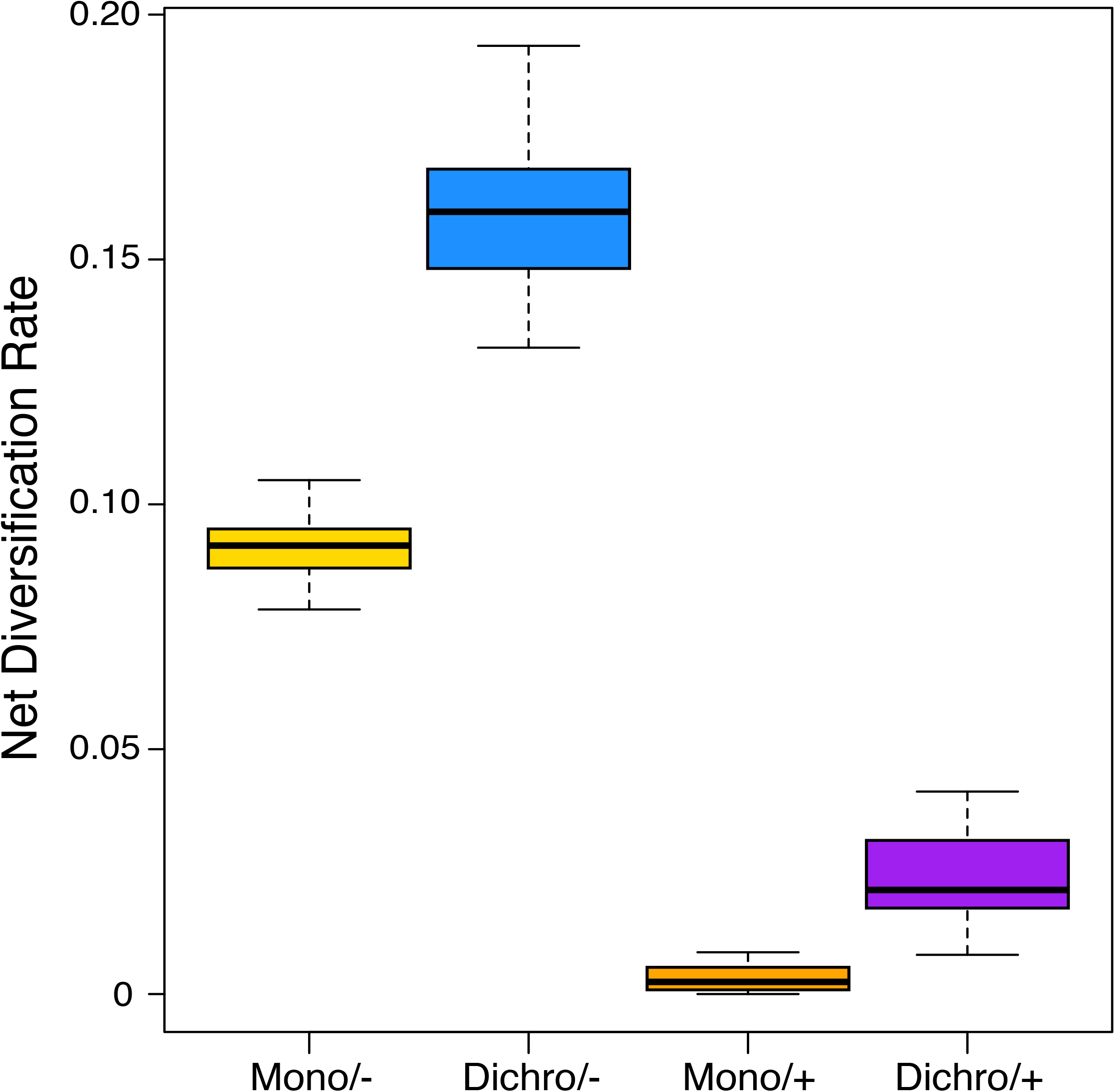
The net diversification rate confidence intervals estimated from the best-fit HiSSE model for various combinations of monochromatism, dichromatism, and the presence (+) or absence (-) of the hidden state.

#### Simulated traits and state-dependent diversification

Analyzed in the BiSSE framework, we found that many of the trait simulations on the phylogeny of Afrobatrachia resulted in the rejection of the ‘null’ model of equal diversification rates across character states. Based on the significance of likelihood ratio tests, we rejected the ‘null’ model in favor of trait-dependent diversification for 565 (37.6%) of our 1500 comparisons. We recovered similar results using a delta AIC cutoff value of two, for which we found support for trait-dependent diversification in 546 (36.4%) of the simulations (Fig. 4). In many of these cases we found unexpectedly strong support for the BiSSE model, and 160 (10.6%) of the comparisons resulted in delta AIC values ranging from 10–40.

**Figure 4.**
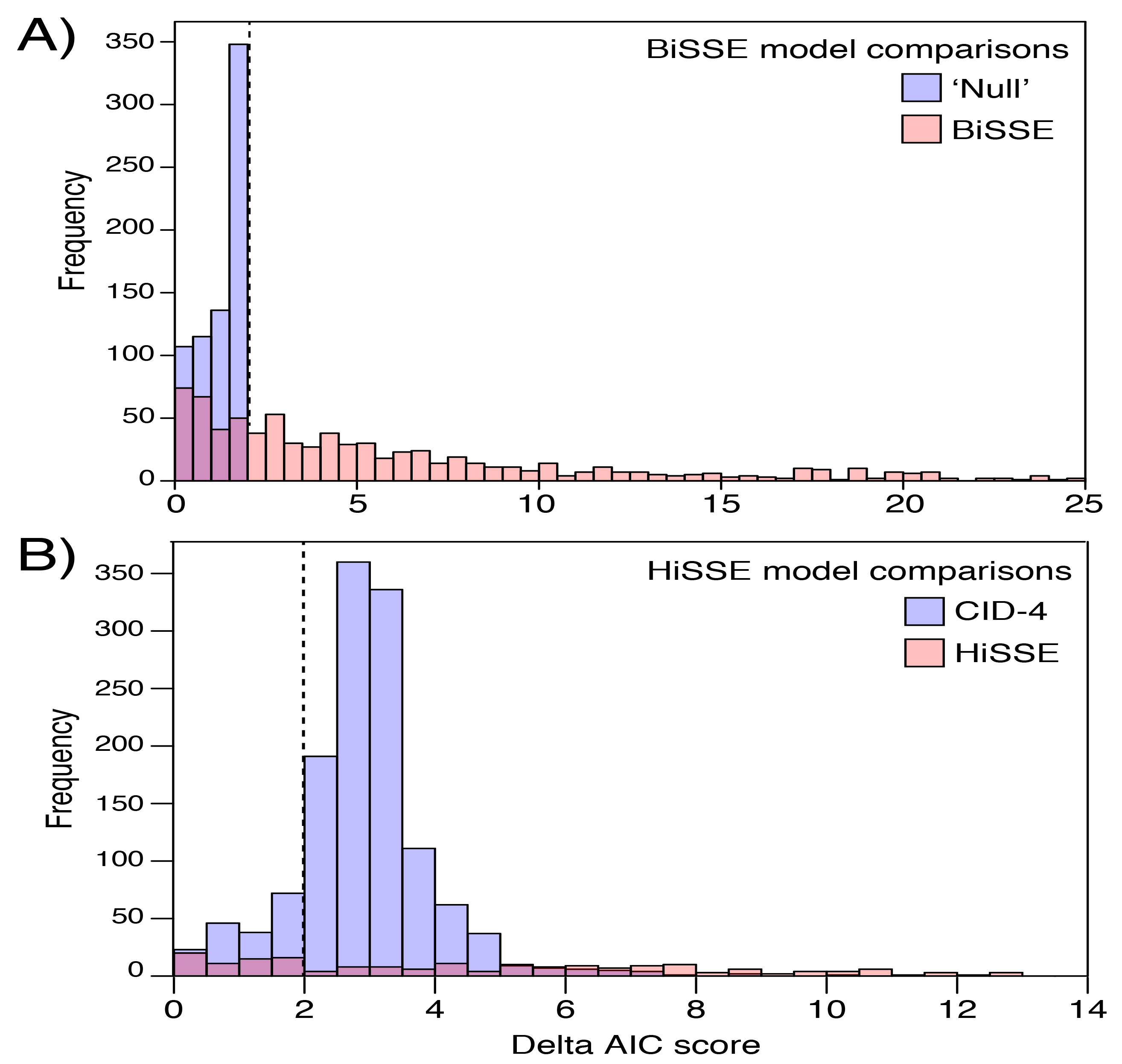
Histograms of the delta AIC comparisons resulting from (A) BiSSE analyses and (B) HiSSE analyses of the 1,500 simulations of neutrally evolving traits on the Afrobatrachia phylogeny. The vertical dotted line represents a delta AIC of two. In the BiSSE analyses (A), delta AIC scores of less than two are considered a failure to reject the ‘null’ model of constant diversification rates, whereas scores above two for the BiSSE model are interpreted as favoring a correlation between diversification rates and the simulated neutral trait. The HiSSE analyses (B) included five models, but only two were consistently selected as the top ranking, including the HiSSE and CID-4 (character-independent diversification) models. For these analyses delta AIC > 2 was considered as support for a particular model, whereas delta AIC < 2 was considered as equivocal support for the model.

In contrast to the BiSSE analyses, the addition of the character independent models (CID-2 and CID-4) in HiSSE dramatically reduced the detection of a false association between simulated traits and diversification rates by providing appropriate null models. In addition to these two CID models, our set of five models also included two BiSSE-like models and a typical HiSSE model. Based on a delta AIC cutoff value of two, out of the 1500 analyses performed the CID-4 model was selected 1,132 times (75.4%), the HiSSE model was chosen 125 times (8.3%), and the remaining 243 analyses (16.2%) showed equivocal support (ΔAIC = 0–2) for either the CID-4 or HiSSE model (Fig. 5). In the cases of equivocal support, the CID-4 and HiSSE models were always the top two models, which should be interpreted as a lack of support for trait-dependent diversification. Our error rate with the HiSSE model being favored in only 8.3% of our simulations was substantially lower than the Beaulieu and O’Meara (2016) ‘difficult tree’ scenario in which the HiSSE model was favored in 29% of the data sets. These simulation results strongly suggest the branching pattern of our empirical phylogeny is not inherently problematic for the investigation of trait-dependent diversification using these available methods, adding support to our empirical analyses in which we detected an association between diversification rates and sexual dichromatism.

**Figure 5.**
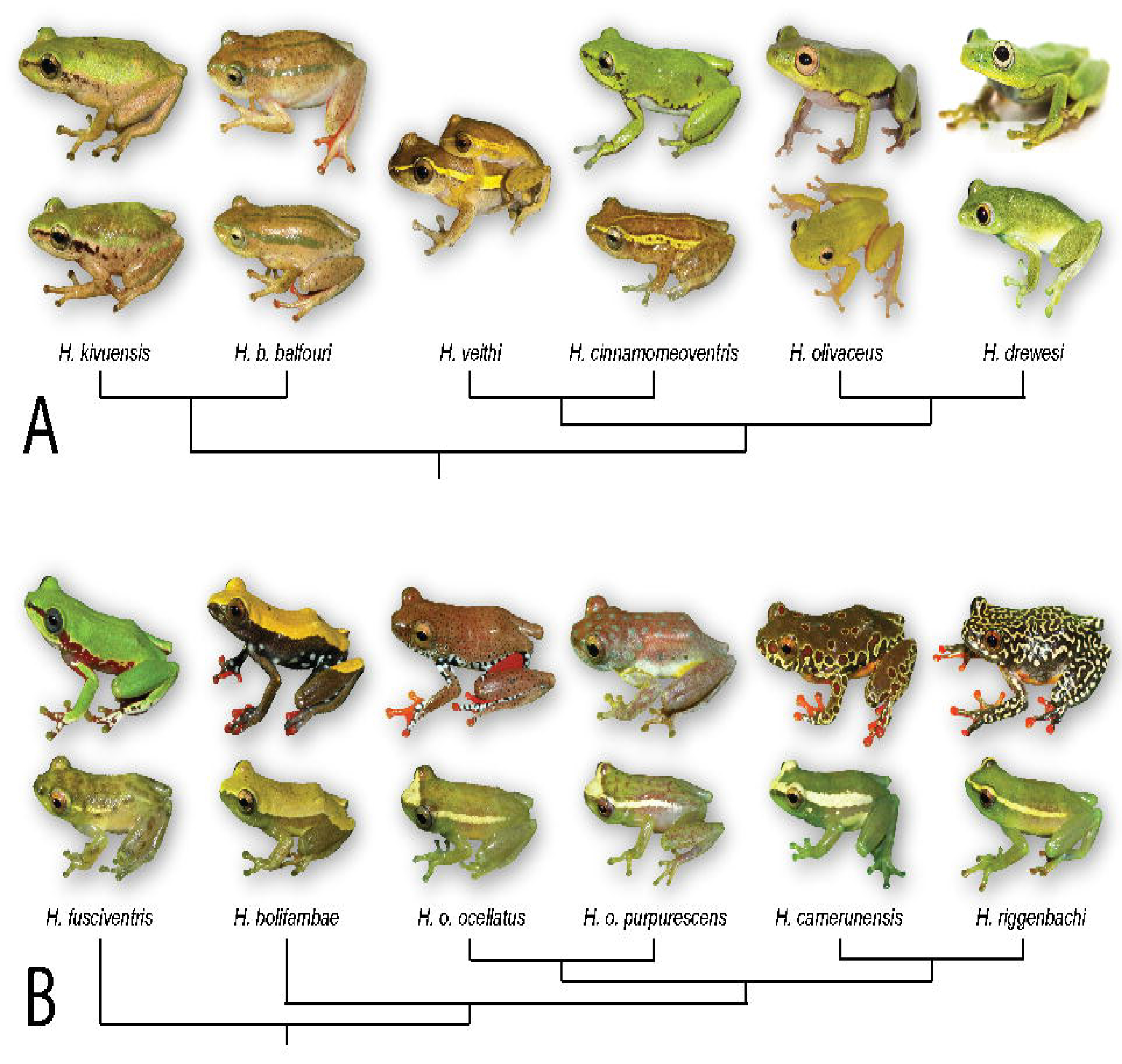
Illustration of (A) several *Hyperolius* species in the predominately sexually dichromatic Clade 1 and (B) several *Hyperolius* species in Clade 2 that exhibit multiple transitions to secondary monochromatism. Females are positioned in the top row, with males below (unless shown in amplexus), and the phylogenetic relationships among species are depicted (though not all species have been included).

## DISCUSSION

### Hyperoliid Relationships and the Origin of Sexual Dichromatism

Afrobatrachian frogs account for more than half of all African amphibians and this continental radiation exhibits incredible variation in ecomorphology, reproductive mode, and other life history traits (Portik & Blackburn 2016). Within Afrobatrachia, the family Hyperoliidae is the most species-rich (~230 species) with surprisingly little diversity in ecomorphology and reproductive mode (Schiøtz 1967, 1999; Portik & Blackburn 2016), but with incredible variation in coloration and sexual dichromatism. These phenotypic characteristics have generated considerable taxonomic confusion within hyperoliids (e.g. Ahl 1931), hindering a clear understanding of species diversity and the evolutionary history of this radiation. Here, we have produced the most comprehensive hyperoliid species tree to date and found support for an Eocene origin of two subfamilies representing a major division withiin Hyperoliidae: Kassininae (26 species) and Hyperoliinae (~200 species). Within Hyperoliinae, we clarified relationships within the hyperdiverse genus *Hyperolius* (~150 species), which largely consists of two clades (Figs. 2, S1, S2). These clades represent parallel rapid radiations with species distributed across a variety of habitats and altitudes throughout sub-Saharan Africa, which showcases Hyperoliidae as a rich comparative framework for future biogeographic research.

While both Kassininae and Hyperoliinae include colorful species, sexual dichromatism only occurs within Hyperoliinae where it is present in the genera *Tachycnemis*, *Cryptothylax* and *Morerella*, several species of *Heterixalus*, and more than half of the *Hyperolius* species we sampled (Fig. 2, Table S2). A previous study hypothesized that sexual dichromatism evolved multiple times within Hyperoliidae (Veith et al. 2009); however, we found overwhelming support for a single origin of sexual dichromatism (Figs. 2, S3) and over twenty independent reversals to monochromatism that range from early transitions that characterize entire genera (e.g., *Afrixalus*) to recent reversals within *Heterixalus* and *Hyperolius* (Figs. 2, S4). This transition bias from dichromatism to monochromatism also occurs in birds (Price & Birch 1996; Omland 1997; Burns 1998; Kimball et al. 2001; Hofmann et al. 2008; Dunn et al. 2015; Schultz & Burns 2017), in which secondary monochromatism can evolve as the result of a color change in either sex (Kimball & Ligon 1999; Johnson et al. 2013; Price & Eaton 2014; Dunn et al. 2015; Schultz et al. 2017). Similar transitional pathways to secondary monochromatism occur in hyperoliids, in which females may lose the ability for color transformation at sexual maturity (both sexes retain the Phase J juvenile coloration) or males undergo obligatory ontogenetic color change at maturity (both sexes develop Phase F coloration) (Figs. 1, 5). Characterizing juvenile coloration across species, which is undocumented in many hyperoliids, and documenting ontogenetic color change, including the prevalence of male color phases (Portik et al. 2016a), will be essential steps toward differentiating between these two forms of monochromatism. In experimental settings, the hormone estradiol induces color transformation in both sexes in the dichromatic species *H. argus* and *H. viridiflavus*, yet the effects of testosterone differ across species (Richards 1982; Hayes & Menendez 1999) suggesting that evolutionary transitions from dichromatism to monochromatism result directly from differences in hormone sensitivities among species. Reversals to monochromatism are especially prominent in *Hyperolius* Clade 2 (twelve independent events; Figs. 2, S4), highlighting the potential for future research in this group to identify the physiological basis and molecular underpinnings of these transitions. For example, three monochromatic species of the *H. cinnamomeoventris* complex that are endemic to the islands of São Tomé and Príncipe (*H. drewesi*, *H. molleri*, *H. thomensis*; both sexes with Phase F coloration) are derived from a mainland clade with sexually dichromatic (*H. cinnamomeoventris*, *H. olivaceus*; females Phase F, males Phase J) and monochromatic species (*H. veithi*; both sexes Phase J) (Fig. 5). Variation among these closely related species is well suited for investigating both the evolutionary and ecological contexts underlying transitions from sexual dichromatism to monochromatism.

### Sexual Dichromatism is Linked to Increased Diversification Rates

Sexual dichromatism is an important predictor of diversification in several taxonomic groups including cichlids (Wagner et al. 2012), labrid fish (Alfaro et al. 2009; Kazancıo⍰lu et al. 2009), agamid lizards (Stuart-Fox & Owens 2003), and dragonflies (Misof 2002); yet in birds, the relationship between dichromatism and species richness/speciation rate varies among studies that differ in methodology and evolutionary scale (Barraclough et al. 1995; Owens et al. 1999; Morrow et al. 2003; Phillimore et al. 2006; Seddon et al. 2013; Huang & Rabosky 2014). Though some state-dependent speciation and extinction model sets, such as the BiSSE method, are no longer considered adequate for robustly detecting trait-dependent diversification (Rabosky & Goldberg 2015; Beaulieu & O’Meara 2016), the HiSSE method contains an expanded model set that can link diversification rate heterogeneity to observed traits, hidden traits, or character independent processes. In particular, the inclusion of appropriate null models (character-independent models) reduces the inference of trait-dependent diversification when a phylogeny contains diversification rate shifts unrelated to the focal trait (Beaulieu & O’Meara 2016). Our trait simulation study recapitulated this result (Fig. 4) while also demonstrating that the branching pattern of our afrobatrachian phylogeny is unlikely to drive false inferences of trait-dependent diversification using HiSSE (e.g., the “worst-case” scenario of Beaulieu & O’Meara 2016). We found that sexually dichromatic hyperoliid lineages have nearly double the average diversification rate of monochromatic lineages, and that these diversification rates are not inflated by a hidden trait. Arboreal oviposition and an associated shift to breeding in pond systems both evolved within hyperoliids and were previously suggested as potential factors influencing diversification (Portik & Blackburn 2016). In our analyses these traits may have represented hidden factors co-distributed with dichromatism; however, the shift to the sexual dichromatism plus hidden state character combination occurred only once in the common ancestor of two genera (*Cryptothylax*, *Morerella*; Fig. S5) and is actually associated with lower diversification rates (Fig. 3), indicating that these additional traits had little impact on our analyses. Together, our results demonstrate that diversification rate heterogeneity occurs within Afrobatrachia and that the origin and persistence of sexual dichromatism in hyperoliid frogs is linked to their rapid diversification across sub-Saharan Africa.

### How Does Sexual Dichromatism Influence Diversification Rate?

Sexual dichromatism is a common proxy for sexual selection in macroevolutionary studies (reviewed in Kraaijeveld et al. 2011), especially for testing the prediction that clades with variation in secondary sexual characters under strong sexual selection exhibit higher diversification rates and species richness (Lande 1981, 1982; West-Eberhard 1983; Barraclough et al. 1995). While our trait-dependent diversification analyses strongly support a faster rate of net diversification associated with sexual dichromatism, we currently lack behavioral and natural history data in hyperoliids to establish whether sexual dichromatism is actually under strong sexual selection. At more recent timescales, speciation by sexual selection can result in a group of closely related species with high ecological similarity but differing almost exclusively in signaling traits (West-Eberhard 1983; Dominey 1984; Panhuis et al. 2001; Turelli et al. 2001; Andersson 2004; Mendelson & Shaw 2005; Ritchie 2007; Safran et al. 2013). Frogs are generally opportunistic gape-limited predators (Duellman & Trueb 1986), and in hyperoliids food partitioning is strongly dictated by body size (Luiselli et al. 2004). In one well-characterized community site, the females of seven syntopic sexually dichromatic *Hyperolius* species display minimal body size differences (Portik et al. 2018) but show exceptional divergence in color (Portik et al. 2016a). Five of these species occur in the predominantly dichromatic *Hyperolius* Clade 1, and within this clade there are typically striking interspecific color differences in females, but not males, among species (Fig. 5). This pattern of interspecific variation in secondary sexual characters in the absence of ecological divergence is consistent with the predictions of speciation by sexual selection, and by inference, would imply that dichromatism in hyperoliids – specifically female color – is an essential mate recognition signal (*sensu* Mendelson & Shaw 2012). In frogs, male calls are well-established mate recognition signals that can be under strong sexual selection (Ryan 1980; Gerhardt 1994), and these acoustic signals are demonstrably important for hyperoliid frogs. Males form dense breeding aggregations and choruses (Bishop et al. 1995), calls differ between closely-related species (Schiøtz 1967, 1999; Gilbert & Bell 2018), and females prefer conspecific calls over calls of syntopic heterospecifics (Telford & Passmore 1981). However, visual displays can also serve as important courtship signals in frogs, typically in conjunction with acoustic signaling (Gomez et al. 2009; Gomez et al. 2010; Starnberger et al. 2014; Jacobs et al. 2016; Yovanovich et al. 2017; Akopyan et al. 2018). Although most studies have documented female preference for male coloration, the recent discovery of female displays during nocturnal phyllomedusine treefrog courtship highlights the possibility of male mate-choice and intersexual selection of female coloration (Akopyan et al. 2018). The lack of heterospecific matings among dichromatic species at high-diversity breeding sites (Portik et al. 2018) strongly suggests behavioral isolation may be linked to divergent mate recognition signals, including male call (Telford & Passmore 1981), gular gland compounds (Starnberger et al. 2013), or female coloration. Our knowledge of reproductive behavior in hyperoliids is largely based on a single dichromatic species (*H. marmoratus*; Dyson & Passmore 1988; Telford & Dyson 1988; Dyson et al. 1992; Jennions et al. 1995), and as such there may be an overlooked role for mutual mate-choice in hyperoliids, in which females locate males by call and/or pheromones and males assess color patterns of approaching females.

Although it can be tempting to equate sexually dimorphic traits such as dichromatism with sexual selection, several alternative mechanisms may also contribute to female-biased dichromatism in hyperoliids. For instance, aposematism is a widespread anti-predator mechanism in frogs that is typically accompanied by the presence of skin toxins (reviewed in Toledo & Haddad 2009; Rojas 2017), commonly alkaloids (Daly 1995). Sex-specific differences in chemical defense occur in some frogs (Saporito et al. 2010; Jeckel et al. 2015); however, both males and females of four dichromatic hyperoliids lacked alkaloids in their skin (Portik et al. 2015) suggesting that either aposematism is an unlikely explanation for ornate coloration or that hyperoliids have evolved novel compounds for chemical defense. Female-biased dichromatism has also been tied to sex-role reversal in fishes and birds (Roede 1972; Oring 1982; Berglund et al. 1986a, 1986b; Eens & Pinxten 2000), in which females compete more intensely than males for access to mates. This mechanism seems unlikely for hyperoliids because males in both monochromatic and dichromatic species form dense choruses and compete intensely to attract females, often engaging in combat (Telford 1985; Backwell & Passmore 1990). Finally, sexual niche partitioning (Selander 1966; Shine 1989) can result in sexual dichromatism when the sexes use different microhabitats, and as a consequence are subject to different selective pressures and predation regimes (Heinsohn et al. 2005; Bell & Zamudio 2012). During the breeding season, male hyperoliids are exposed on calling sites and Hayes (1997) proposed that more cryptic male coloration may reduce predation pressure, a hypothesis that is consistent with greater observations of predation events on females of dichromatic species (Grafe 1997; Portik et al. 2018). Quantifying differences in predation rates between the sexes, between monochromatic and dichromatic species, and between Phase F and Phase J males within dichromatic species would address whether this aspect of natural selection is also shaping the evolution of sexual dichromatism. Although these mechanisms and other differences in natural selection pressures may influence the evolution of sexual dichromatism in hyperoliids, they are generally not expected to elevate rates of reproductive isolation or drive diversification rate shifts comparable to the effects of sexual selection. Therefore, we propose that hyperoliids are a compelling system for disentangling the roles of sexual selection and natural selection in the evolution of sexual dichromatism, and how these mechanisms have promoted diversification at both microevolutionary and macroevolutionary timescales.

## ACKNOWLEDGEMENTS

Laboratory work conducted by DMP was funded by a National Science Foundation DDIG (DEB: 1311006), a National Science Foundation grant (DEB: 1202609) awarded to D.C. Blackburn, an Ecological, Evolutionary, and Conservation Genomics Research Award presented by the American Genetic Association, the Museum of Vertebrate Zoology, and by R.C. Bell and J.A. McGuire. This work used the Vincent J. Coates Genomics Sequencing Laboratory at UC Berkeley, supported by NIH S10 Instrumentation Grants S10RR029668 and S10RR027303. We thank the following institutions for accessioning field collections and for facilitating loan access: California Academy of Sciences, Cornell University Museum of Vertebrates, Institut national de Recherche en Sciences Exactes et Naturelles, Museum für Naturkunde, Berlin, Muséum National d’Histoire Naturelle, Museum of Comparative Zoology, Museum of Vertebrate Zoology, National Museum, Prague, Natural History Museum, London, North Carolina Museum of Natural Sciences, Port Elizabeth Museum, Senckenberg Natural History Collections Dresden, South African Institute for Aquatic Biodiversity, South African National Biodiversity Institute, The Field Museum, Trento Museum of Science, University of Texas at El Paso Biodiversity Collections, and the Zoological Natural History Museum, Addis Ababa University. All authors express thanks to the many government agencies, ministries, and departments that issued research permits and provided the access necessary to conduct their individual field work in sub-Saharan Africa.

## SUPPLEMENTAL MATERIAL

Figure S1. A species tree of Hyperoliidae inferred from 1,047 gene trees with ASTRAL-III. Node support is shown for quartet support values and multi-locus bootstraps, with branch lengths depicted in coalescent units (except for tree tips, represented by dotted lines).

Figure S2. A chronogram of Afrobatrachia inferred from the multilocus BEAST analysis of 283 species. Taxa labeled with blue color denote the 153 species included in the hyperoliid species tree that was used as a partial constraint tree in this analysis. Node support values are not shown because the tree topology was fixed, but error bars representing the 95% HPD for dating estimates are provided.

Figure S3. A comparison of Bayesian ancestral character reconstructions inferred using the stochastic Mk model of character evolution with symmetrical (Mk1) or asymmetrical (Mk2) transition rates in combination with a strict clock or random local clock model. All analyses enforced a monochromatic root prior. The marginal likelihoods and average number of state changes are shown for each analysis. Node sizes reflect the posterior probability for the inferred character state, and tips and nodes are colored with monochromatism in yellow and dichromatism in blue.

Figure S4. An illustration of transitions to secondary monochromatism inferred using Bayesian ancestral character reconstruction with the Mk2 and strict clock models. Large nodes outlined with red represent inferred transitions to monochromatism from a dichromatic ancestor. States at nodes and tips are colored with monochromatism in yellow and dichromatism in blue.

Figure S5. Ancestral state reconstruction of the observed and hidden state combinations in Afrobatrachian frogs from the best-fit HiSSE model (HiSSE 19; Table 1). Squares at tips are colored by the observed character states (yellow: monochromatic, blue: sexually dichromatic), whereas node pie charts represent the probability of a state assignment to: 1) monochromatism + hidden state absent (yellow), 2) monochromatism + hidden state present (orange), 3) dichromatism + hidden state absent (blue), or 4) dichromatism + hidden state present (purple). No instance of the monochromatism + hidden state present is reconstructed on the phylogeny, whereas a transition to the dichromatism + hidden state present combination is inferred once in the MRCA of two monotypic genera (*Cryptothylax*, *Morerella*), indicating a near absence of the hidden state across the entire tree.

Table S1. Taxon, voucher, and locality information for all samples included in the sequence-capture experiment.

Table S2. A summary of the occurrence of monochromatism and sexual dichromatism in Afrobatrachian species based on literature, museum specimens, and collective field observations.

## LITERATURE CITED

Ahl E (1931) *Amphibia, Anura III, Polypedatidae*. Das Tierreich 55: xvi + 477.

Akopyan M, Kaiser K, Vega A, Savant NG, Owen CY, Dudgeon SR, Robertson JM (2018) Melodic males and flashy females: geographic variation in male and female reproductive behavior in red-eyed treefrogs (*Agalychnis callidryas*). Ethology 124:54−64.

Alfaro ME, Brock CD, Banbury BL, Wainwright PC (2009) Does evolutionary innovation in pharyngeal jaws lead to rapid lineage diversification in labrid fishes? BMC Evol Biol 9:255.

Aljanabi S, Martinez I (1997) Universal and rapid salt-extraction of high quality genomic DNA for PCR-based techniques. Nucleic Acids Res 25:4692–4693.

Amiet J-L (2012) Les Rainettes du Cameroun (Amphibiens Anoures). Saint-Nazaire, France: La Nef des Livres. 591 pp.

AmphibiaWeb (2018) Available at: amphibiaweb.org. University of California, Berkeley. Accessed May 2018.

Andersson M (1994) Sexual selection. Princeton, NJ. Princeton University Press.

Backwell PRY, Passmore NI (1990) Aggressive interactions and intermale spacing in choruses of the leaf-folding frog, *Afrixalus delicatus*. S Afr J Zool 25:133–137.

Baele G, Lemey P, Bedford T, Rambaut A, Suchard MA, Alekseyenko AV (2012) Improving the accuracy of demographic and molecular clock model comparison while accommodating phylogenetic uncertainty. Mol Biol Evol 29:2157–2167.

Baele G, Li WLS, Drummond AJ, Suchard MA, Lemey P (2012) Accurate model selection of relaxed molecular clocks in Bayesian phylogenetics. Mol Biol Evol 30:239–243.

Barraclough TG, Harvey PH, Nee S (1995) Sexual selection and taxonomic diversity in passerine birds. Proc R Soc Lon B 259:211−215.

Beaulieu JM, O’Meara BC (2016) Detecting hidden diversification shifts in models of trait-dependent speciation and extinction. Syst Biol 65:583−601.

Bell RC, Zamudio KR (2012) Sexual dichromatism in frog: natural selection, sexual selection and unexpected diversity. Proc R Soc Lon B 279:4687−4693.

Bell RC, Parra JL, Badjedjea G, Barej MF, Blackburn DC, Burger M, Channing A, Dehling JM, Greenbaum E, Gvoždík V, Kielgast J, Kusamba C, Lötters S, McLaughlin PJ, Nagy ZT, Rödel M-O, Portik DM, Stuart BL, VanDerWal J, Zamudio KR (2017a) Idiosyncratic responses to climate-driven forest fragmentation and marine incursions in reed frogs from Central Africa and the Gulf of Guinea Islands. Mol Ecol 26:5223–5224.

Bell RC, Webster GN, Whiting MJ (2017b) Breeding biology and the evolution of dynamic sexual dichromatism in frogs. J. Evol. Biol. 30:2104−2115.

Berglund A, Rosenqvist G, Svensson I (1986a) Reversed sex roles and parental energy investment in zygotes of two pipefish (Syngnathidae) species. Mar Ecol Prog Ser 29:209–215.

Berglund A, Rosenqvist G, Svensson I (1986b) Mate choice, fecundity and sexual dimorphism in two pipefish species (Syngnathidae). Behav Ecol Sociobiol 19:301–307.

Bi K, Vanderpool D, Singhal S, Linderoth T, Moritz C, Good JM (2012) Transcriptome-based exon capture enables highly cost-effective comparative genomic data collection at moderate evolutionary scales. BMC Genomics 13:403.

Bishop PJ, Jennions MD, Passmore NI (1995) Chorus size and call intensity: female choice in the painted reed frog, *Hyperolius marmoratus*. Behaviour 132:721–731.

Burnham KP, Anderson DR (2002) Model Selection and Multimodel Inference: A Practical Information-Theoretic Approach. New York: Springer.

Burns K (1998) A phylogenetic perspective on the evolution of sexual dichromatism in tanagers (Thraupidae): the role of female versus male plumage. Evolution 52:1219–1224.

Capella-Gutierrez S, Silla-Martinez JM, Gabaldon T (2009) trimAl: a tool for automated alignment trimming in large-scale phylogenetic analyses. Bioinformatics 25:1972–1973.

Channing A (2001) Amphibians of Central and Southern Africa. Cornell University Press, Ithaca, New York, USA.

Channing A, Howell KM (2006) Amphibians of East Africa. Edition Chimaira, Frankfurt, Germany.

Channing A, Hillers A, Lötters S, Rödel M-O, Schick S, Conradie W, Rödder D, Mercurio V, Wagner P, Dehling JM, du Preez LH, Kielgast J, Burger M (2013) Taxonomy of the super-cryptic *Hyperolius nasutus* group of long reed frogs of Africa (Anura: Hyperoliidae), with descriptions of six new species. Zootaxa 3620:301–350.

Conradie W, Branch WR, Measey GJ, Tolley KA (2012) A new species of *Hyperolius* Rapp, 1842 (Anura: Hyperoliidae) from the Serra da Chela mountains, south-western Angola. Zootaxa 3269:1–17.

Conradie W, Branch WR, Tolley KA (2013) Fifty shades of grey: giving colour to the poorly known Angolan Ashy reed frog (Hyperoliidae: *Hyperolius cinereus*), with the description of a new species. Zootaxa 3635:201–223.

Conradie W, Verbugt L, Portik DM, Ohler A, Bwong BA, Lawson LP (2018) A new reed frog (*Hyperoliidae: Hyperolius*) from coastal northeastern Mozambique. Zootaxa 4379:177−198.

Daly JW (1995) The chemistry of poisons in amphibian skin. Chemical ecology: the chemistry of biotic interaction, eds Eisner T, Meinwald J (National Academy Press, Washington, DC, USA), pp 17–28.

Darwin CR (1871) The Descent of Man, and Selection in Relation to Sex. London: John Murray. Volume 1. 1st edition.

De Lisle, SP, Rowe L (2013) Correlated evolution of allometry and sexual dimorphism across higher taxa. Am Nat 183:630–639.

Dehling JM (2012) An African glass frog: a new *Hyperolius* species (Anura: Hyperoliidae) from Nyungwe National Park, southern Rwanda. Zootaxa 3391:52–64.

Dominey W (1984) Effects of sexual selection and life history on speciation: species flocks in African cichlids and Hawaiian *Drosophila*. Evolution of Fish Species Flocks, eds Echelle AA, Kornfield I (University of Maine Press, Orono), pp 231−249.

Drewes RC, Vindum JV (1994) Amphibians of the Impenetrable Forest, Southwest Uganda. J Afr Zool 108:55–70.

Drummond AJ, Suchard MA, Xie D, Rambaut A (2012) Bayesian phylogenetics with BEAUti and the BEAST 1.7. Mol Biol Evol 29:1969–1973.

Duellman WE, Trueb L (1986) Biology of Amphibians. McGraw-Hill. New York, USA.

Dunn PO, Armenta JK, Whittingham LA (2015) Natural and sexual selection act on different axes of variation in avian plumage color. Sci Adv 1:e1400155.

Dyson ML, Passmore NI (1988) Two-choice phonotaxis in *Hyperolius marmoratus* (Anura: Hyperoliidae): the effect of temporal variation in presented stimuli. Anim Behav 36:648–652.

Dyson ML, Passmore NI, Bishop PJ, Henzi SP (1992) Male behavior and correlates of mating success in a natural population of African painted reed frogs (*Hyperolius marmoratus*). Herpetologica 48:236–246.

Edgar RC (2004) MUSCLE: multiple sequence alignment with high accuracy and high throughput. Nucl Acids Res 32:1792–1797.

Eens M, Pinxten R (2000) Sex-role reversal in vertebrates: behavioural and endocrinological accounts. Behav Process 51:135−147.

Feng Y-J, Blacburn DC, Liang D, Hillis DM, Wake DB, Cannatella DC, Zhang P (2017) Phylogenomics reveals rapid, simultaneous diversification of three major clades of Gondwanan frogs at the Cretaceous-Paleogene boundary. Proc Natl Acad Sci USA 114:E5864−E5870.

FitzJohn RG (2012) Diversitree: comparative phylogenetic analyses of diversification in R. Methods Ecol Evol 3:1084–1092.

FitzJohn RG, Maddison WP, and Otto SP (2009) Estimating trait-dependent speciation and extinction rates from incompletely resolved phylogenies. Syst Biol 58:595−611.

Gerhardt HC (1994) The evolution of vocalization in frogs and toads. Annu Rev Ecol Syst 25:233−324.

Gerhardt HC, Huber F (2002) Acoustic communication in insects and anurans: common problems and diverse solutions. Chicago (IL): The University of Chicago Press.

Gilbert CM, Bell RC (2018) Evolution of advertisement calls in an island radiation of African reed frogs. Biol J Linn Soc 123:1−11.

Gomez D, Richardson C, Lengagne T, Plenet S, Joly P, Léna J-P, Théry M (2009) The role of nocturnal vision in mate choice: females prefer conspicuous males in the European tree frog (*Hyla arborea*). Proc R Soc B 276:2351−2358.

Gomez D, Richardson C, Lengagne T, Derex M, Plenet S, Joly P, Léna J-P, Théry M (2010) Support for a role of colour vision in mate choice in the nocturnal European treefrog. Behaviour 147:1753−1768.

Grafe TU (1997) Costs and benefits of mate choice in the lek-breeding reed frog, *Hyperolius marmoratus*. Anim Behav 53:1103–1117.

Greenbaum E, Sinsch U, Lehr E, Valdez F, Kusamba C (2013) Phylogeography of the reed frog *Hyperolius castaneus* (Anura: Hyperoliidae) from the Albertine Rift of Central Africa: implications for taxonomy, biogeography and conservation. Zootaxa 3131:473–494.

Han X, Fu J (2013) Does life history shape sexual size dimorphism in anurans? A comparative analysis. BMC Evol Biol 13:27.

Hayes TB (1997) Hormonal mechanisms as potential constraints on evolution: examples from the Anura. Am Zool 37:482−490.

Hayes TB, Menendez KP (1999) The effect of sex steroids on primary and secondary sex differentiation in the sexually dichromatic reedfrog (*Hyperolius argus*: Hyperolidae) from the Arabuko Sokoke forest of Kenya. Gen Comp Endocr 115:188−199.

Heinsohn R, Legge S, Endler JA (2005) Extreme reversed sexual dichromatism in a bird without sex role reversal. Science 309:617−619.

Hofmann CM, Cronin TW, Omland KE (2008) Evolution of sexual dichromatism. 1. Convergent losses of elaborate female coloration in New World orioles (*Icterus* spp.) Auk 125:778−789.

Huang H, Rabosky DL (2014) Sexual selection and diversification: reexamining the correlation between dichromatism and speciation rate in birds. Am Nat 184(5):E104−E114.

Jacobs LE, Vega A, Dudgeon S, Kaiser K, Robertson JM (2016) Local not vocal: assortative female choice in divergent populations of red-eyed treefrogs, *Agalychnis callidryas* (Hylidae: Phyllomedusinae). Biol J Linn Soc 120:171−178.

Jeckel AM, Grant T, Saporito RA (2015) Sequestered and synthesized chemical defenses in the poison frog *Melanophryniscus moreirae*. J Chem Ecol 41:505–512.

Jennions MD, Bishop PJ, Backwell PRY, Passmore NI (1995) Call rate variability and female choice in the African frog, *Hyperolius marmoratus*. Behaviour 132:709–720.

Johnson AE, Price JJ, Pruett-Jones S (2013) Different modes of evolution in males and females generate dichromatism in fairy-wrens (Maluridae). Ecol Evol 3:3030–3046.

Kazancıo⍰lu E, Near TJ, Hanel R, Wainwright PC (2009) Influence of sexual selection and feeding functional morphology on diversification rate of parrotfishes. Proc R Soc B 276:3439−3446.

Katoh K, Standley DM (2013) MAFFT multiple sequence alignment software version 7: improvements in performance and usability. Mol Biol Evol 30:722–780.

Katoh K, Kuma K, Toh H, Miyata T (2005) MAFFT version 5: improvement in accuracy of multiple sequence alignment. Nucleic Acids Res 33:511–518.

Katoh K, Misawa K, Kuma K, Miyata T (2002) MAFFT: a novel method for rapid multiple sequence alignment based on fast Fourier transform. Nucleic Acids Res 30:3059–3066.

Kimball RT, Ligon JD (1999) Evolution of avian plumage dichromatism from a proximate perspective. Am Nat 154(2):182−193.

Kimball RT, Braun EL, Ligon JD, Lucchini V, Randi E (2001) A molecular phylogeny of the peacock-pheasants (Galliformes: *Polyplectron* spp.) indicates loss and reduction of ornamental traits and display behaviours. Biol J Linn Soc 73:187−198.

Kindermann C, Hero J-M (2016) Rapid dynamic colour change is an intrasexual signal in a lek breeding frog (*Litoria wilcoxii*). Behav Ecol Sociobiol 70:1995−2003.

King B, Lee MSY (2015) Ancestral state reconstruction, rate heterogeneity, and the evolution of reptile viviparity. Syst Biol 64:532−544.

Kirkpatrick M (1982) Sexual selection and the evolution of female choice. Evolution 36:1−12.

Kouamé N’GG, Boateng CO, Rödel M-O (2013) A rapid survey of the amphibians from the Atewa Range Forest Reserve, Eastern Region, Ghana. A Rapid Biological Assessment of the Atewa Range Forest Reserve, Eastern Ghana, ed. Conservation International (Arlington, Virginia), pp 76–83.

Kouamé AM, Kouamé N’GG, Konan JCBYN’G, Adepo-Gourène B, Rödel M-O (2015) Contributions to the reproductive biology and behaviour of the dotted reed frog, *Hyperolius guttulatus*, in southern-central Ivory Coast, West Africa. Herpetology Notes 8:633−641.

Kraaijeveld K, Kraaijeveld-Smit FJL, Maan ME (2011) Sexual selection and speciation: the comparative evidence revisited. Biol Rev 86:367−377.

Kurabayashi A, Sumida M (2013) Afrobatrachian mitochondrial genomes: genome reorganization, gene rearrangement mechanisms, and evolutionary trends of duplicated and rearranged genes. BMC Genomics 14:633.

Lande R (1981) Models of speciation by sexual selection on polygenic traits. Proc Natl Acad Sci USA 78:3721−3725.

Lande R (1982) Rapid origin of sexual isolation and character displacement in a cline. Evolution 36:213−223.

Lewis PO (2001) A likelihood approach to estimating phylogeny from discrete morphological character data. Syst Biol 50:913–925.

Liedtke HC, Hügil D, Dehling JM, Pupin F, Menegon M, Plumptre AJ, Kujirakwinja D, Loader SP (2014) One or two species? On the case of *Hyperolius discodactylus* Ahl, 1931 and *H. alticola* Ahl, 1931 (Anura: Hyperoliidae). Zootaxa 3768:253–290.

Liu L, Yu L, Edwards SV (2010) A maximum pseudo-likelihood approach for estimating species trees under the coalescent model. BMC Evol Biol 10:302.

Loader SP, Ceccarelli FS, Menegon M, Howell KM, Kassahun R, Mengistu AA, Saber SA, Gebresenbet F, de Sa R, Davenport TRB, Larson JG, Müller H, Wilkinson M, Gower DJ (2014) Persistence and stability of Eastern Afromontane forests: evidence from brevicipitid frogs. J Biogeogr 41:1781–1792.

Loader SP, Lawson LP, Portik DM, Menegon M (2015) Three new species of spiny throated reed frogs (Anura: Hyperoliidae) from evergreen forests of Tanzania. BMC Research Notes 8:167.

Luiselli L, Bikikoro L, Odegbune E, Wariboko SM, Rugiero L, Akani GC, Politano E (2004) Feeding relationships between sympatric Afrotropical frogs (genus *Hyperolius*): the effects of predator body size and season. Animal Biology 54:293−302.

Maddison WP, Midford PE, Otto SP (2007) Estimating a binary character’s effect on speciation and extinction. Syst Biol 56:701−710.

Maan ME, Cummings ME (2009) Sexual dimorphism and directional sexual selection on aposematic signals in a poison frog. Proc Natl Acad Sci USA 106:19072–19077.

McDiarmid RW (1975) Glass frog romance along a tropical stream. Nat. Hist. Mus. Los Angeles Co. Terra. 13(4):14−18.

Mendelson TC, Shaw KL (2012) The (mis)concept of species recognition. Trends Ecol. Evol. 27:421−427.

Meyer M, Kircher M (2010) Illumina sequencing library preparation for highly multiplexed target capture and sequencing. Cold Spring Harbor Protocols 2010:pdb.prot5448.

Mirarab S, Warnow T (2015) ASTRAL-II: coalescent-based species tree estimation with many hundreds of taxa and thousands of genes. Bioinformatics 31:i44–i52.

Mirarab S, Reaz R, Bayzid MdS., Zimmermann T, Swenson MS, Warnow T (2014) ASTRAL: genome-scale coalescent-based species tree estimation. Bioinformatics 30:i541–i548.

Misof B (2002) Diversity of Anisoptera (Odonata): inferring speciation processes from patterns of morphological diversity. Zoology 105:355−365.

Morrow EH, Pitcher TE, Anrqvist G (2003) No evidence that sexual selection is an ‘engine of speciation’ in birds. Ecol Lett 6:228−234.

Nali RC, Zamudio KR, Haddad CFB, Prado CPA. 2014. Size-dependent selective mechanisms on males and females and the evolution of sexual size dimorphism in frogs. Am Nat 184:727–740.

Omland KE (1997) Examining two standard assumptions of ancestral reconstructions: repeated loss of dichromatism in dabbling ducks. Evolution 51:1636−1646.

Oring LW (1982) Avian mating systems. Avian Biology, eds Farner DS, King JS, Parkes KC (Academic Press, New York), pp 1–92.

Owens IPF, Bennett PM, Harvey PH (1999) Species richness among birds: body size, life history, sexual selection or ecology? Proc R Soc Lon B 266:933−939.

Palumbi S, Martin A, Romano S, McMillan WO, Stice L, Grabowski G (1991) The Simple Fool’s Guide to PCR. Version 2. Honolulu: University of Hawaii.

Panhuis TM, Butlin R, Zuk M, Tregenza T (2001) Sexual selection and speciation. Trends Ecol Evol 16(7):364−371.

Paradis E, Claude J, Strimmer K (2004) APE: analysis of phylogenetics and evolution in R language. Bioinformatics 20:289–290.

Phillimore AB, Freckleton RP, Orme CDL, Owens IPF (2006) Ecology predicts large-scale patterns of phylogenetic diversification in birds. Am Nat 168(2):220−229.

Portik DM (2015. Diversification of Afrobatrachian frogs and the herpetofauna of the Arabian Peninsula. PhD Thesis. University of California, Berkeley.

Portik DM, Blackburn DC (2016) The evolution of reproductive diversity in Afrobatrachia: a phylogenetic comparative analysis of an extensive radiation of African frogs. Evolution 70(9):2017−2032.

Portik DM, Scheinberg L, Blackburn DC, Saporito RA (2015) Lack of defensive alkaloids in the integumentary tissue of four brilliantly colored African reed frog species (Hyperoliidae. *Hyperolius*). Herpetological Conservation and Biology 10:833−838.

Portik DM, Jongsma GF, Kouete MT, Scheinberg LA, Freiermuth B, Tapondjou WP, Blackburn DC (2016a) A survey of amphibians and reptiles in the foothills of Mount Kupe, Cameroon. Amphibian and Reptile Conservation 10(2)[Special Section]:37–67(e131).

Portik DM, Smith LL, Bi K (2016b) An evaluation of transcriptome-based exon capture for frog phylogenomics across multiple scales of divergence (Class: Amphibia, Order: Anura). Mol Ecol Resour 16:1069–1083.

Portik DM, Smith LL, Bi K (2016c) Data from: An evaluation of transcriptome-based exon capture for frog phylogenomics across multiple scales of divergence (Class: Amphibia, Order: Anura). Dryad Digital Repository. https://doi.org/10.5061/dryad.pr3pr.

Portik DM, Jongsma GF, Kouete MT, Scheinberg LA, Freiermuth B, Tapondjou WP, Blackburn DC (2018). Ecological, morphological, and reproductive aspects of a diverse assemblage of hyperoliid frogs (Family: Hyperoliidae) surrounding Mt. Kupe, Cameroon. Herpetol Rev, In Press.

Price JJ, Eaton MD (2014) Reconstructing the evolution of sexual dichromatism: current color diversity does not reflect past rates of male and female change. Evolution 68:2026−2037.

Price T (1998) Sexual selection and natural selection in bird speciation. Phil Trans R Soc Lond B 353:251−260.

Price T, Birch GL (1996) Repeated evolution of sexual color dimorphism in passerine birds. Auk 113:842−848.

Pyron RA, Wiens JJ. 2011. A large-scale phylogeny of Amphibia including over 2800 species, and a revised classification of extant frogs, salamanders, and caecilians. Mol Phylogenet Evol 61:543–583.

Rabosky DL, Goldberg EE (2015) Model inadequacy and mistaken inferences of trait-dependent speciation. Syst Biol 64:340–355.

Rambaut A, Drummond AJ, Suchard M (2013). Tracer v1.6.0. Available from: http://beast.bio.ed.ac.uk/

Revell LJ (2012) Phytools: an R package for phylogenetic comparative biology (and other things). Methods Ecol Evol 3:217–223.

Richards CM (1982) The alteration of chromatophore expression by sex hormones in the Kenyan Reed Frog, *Hyperolius viridiflavus*. General and Comparative Endocrinology 46:59−67.

Ritchie MG (2007) Sexual selection and speciation. Ann Rev Ecol Evol S 38:79−102.

Rödel M-O, Grafe TU, Rudolf VHW, Ernst R (2002) A review of West African spotted *Kassina*, including a description of *Kassina schioetzi* sp. nov. (Amphibia: Anura: Hyperoliidae). Copeia 2002:800–814.

Rödel M-O, Kosuch J, Veith M, Ernst R (2003) First record of the genus *Acanthixalus* Laurent, 1944 from the Upper Guinean Rain Forest, West Africa, with the description of a new species. J Herpetol 37:43–52.

Rödel M-O, Kosuch J, Grafe TU, Boistel R, Assemian NE, Kouamé NG, Tohé B, Gourène G, Perret J-L, Henle K, Tafforeau P, Pollet N, Veith M (2009) A new tree-frog genus and species from Ivory Coast, West Africa (Amphibia: Anura: Hyperoliidae). Zootaxa 2044:23–45.

Rödel M-O, Sandberger L, Penner J, Mané Y, Hillers A (2010) The taxonomic status of *Hyperolius spatzi* Ahl, 1931 and *Hyperolius nitidulus* Peters, 1875 (Amphibia: Anura: Hyperoliidae). Bonn Zoological Bulletin 57:177–188.

Roede M (1972) Color as related to size, sex and behavior in seven Caribbean labrid fish species (genera *Thalasoma, Halichoeres* and *Hemipteronotus*). Studies on the fauna of Curacao and other Caribbean Islands 42:1−266.

Roelants K, Gower DJ, Wilkinson M, Loader SP, Biju SD, Guillaume K, Moriau L, Bossuyt F (2007) Global patterns of diversification in the history of modern amphibians. Proc Natl Acad Sci USA 104:887–892.

Ryan MJ (1980) Female choice in a Neotropical frog. Science 209:523−525.

Safran RJ, Scordato ESC, Symes LB, Rodríguez RL, Mendelson TC (2013) Contributions of natural and sexual selection to the evolution of premating reproductive isolation: a research agenda. Trends Ecol. Evol. 28:643−650.

Salthe SN, Duellman WE (1973) Quantitative constraints associated with reproductive mode in anurans. Evolutionary Biology of the Anurans, ed Vial JL (University of Missouri Press, Missouri), pp 229–249.

Sanderson MJ (2002) Estimating absolute rates of molecular evolution and divergence times: a penalized likelihood approach. Mol Biol Evol 19:101–109.

Saporito RA, Donnelly MA, Spande TF, Garraffo HM (2012) A review of chemical ecology in poison frogs. Chemoecology 22:159–168.

Sayyari E, Mirarab S (2016) Fast coalescent-based computation of local branch support from quartet frequencies. Mol Biol Evol 33:1654–1668.

Schick S, Kielgast J, Rödder D, Muchai V, Burger M, Lötters S (2010) New species of reed frog from the Congo Basin with discussion of paraphyly in cinnamon-belly reed frogs. Zootaxa 2501:23–36.

Schiøtz A (1967) The treefrogs (Rhacophoridae) of West Africa. Spolia Zoologica Musei Hauniensis 25:1−346.

Schiøtz A (1999) Treefrogs of Africa. Edition Chimaira, Frankfurt, Germany.

Seddon N, Botero CA, Tobias JA, Dunn PO, MacGregor HEA, Rubenstein DR, Uy AC, Weir JT, Whittingham LA, Safran RJ (2013) Sexual selection accelerates signal evolution during speciation in birds. Proc R Soc B 280:20131065.

Selander RK (1966) Sexual dimorphism and differential niche utilization in birds. The Condor 68:113−151.

Seo T-K (2008) Calculating bootstrap probabilities of phylogeny using multilocus sequence data. Mol Biol Evol 25:960–971.

Shine R (1979) Sexual selection and sexual dimorphism in the Amphibia. Copeia 1979:297–306.

Shine R (1989) Ecological causes for the evolution of sexual dimorphism: a review of the evidence. Q. Rev. Biol. 64(4):419−461.

Shultz AJ, Burns KJ (2017) The role of sexual and natural selection in shaping patterns of sexual dichromatism in the largest family of songbirds (Aves: Thraupidae). Evolution 71:1061−1074.

Singhal S (2013) De novo transcriptomic analyses for non-model organisms: an evaluation of methods across a multi-species data set. Mol Ecol Resour 13:403–416.

Stamatakis A (2014) RAxML version 8: a tool for phylogenetic analysis and post-analysis of large phylogenies. Bioinformatics 30:1312–1313.

Starnberger I, Poth D, Peram PS, Schulz S, Vences M, Knudsen J, Barej MF, Rödel M-O, Walzl M, Hödl W (2013) Take time to smell the frogs: vocal sac glands of reed frogs (Anura: Hyperoliidae) contain species-specific chemical cocktails. Biol J Linn Soc 110:828–838.

Starnberger I, Preininger D, Hödl W (2014) From uni- to multimodality: towards an integrative view on anuran communication. J Comp Physiol A 200:777−787.

Stuart-Fox D, Owens IPF (2003) Species richness in agamid lizards: chance, body size, sexual selection or ecology? J Evolution Biol 16:659−669.

Sztatecsny M, Preininger D, Freudmann A, Loretto M-C, Maier F, Hödl W (2012) Don’t get the blues: conspicuous nuptial colouration of male moor frogs (*Rana arvalis*) supports visual mate recognition during scramble competition in large breeding aggregations. Behav Ecol Sociobiol 66:1587−1593.

Telford SR, Dyson ML (1988) Some determinants of the mating system in a population of painted reed frogs (*Hyperolius marmoratus*). Behaviour 106:265–278.

Telford SR, Passmore NI (1981) Selective phonotaxis of four sympatric species of African reed frogs (genus *Hyperolius*). Herpetologica 37:29–32.

Turelli M, Barton NH, Coyne JA (2001) Theory and speciation. Trends Ecol Evol 16(7):330−342.

Veith M, Kosuch J, Rödel M-O, Hillers A, Schmitz A, Burger M, Lötters S (2009) Multiple evolution of sexual dichromatism in African reed frogs. Mol Phylogenet Evol 51:388−393.

Wagner CE, Harmon LJ, Seehausen O (2012) Ecological opportunity and sexual selection together predict adaptive radiation. Nature 487:366−369.

Wells KD (1977) The social behavior of anuran amphibians. Anim Behav 25:666–693.

West-Eberhard MJ (1983) Sexual selection, social competition, and speciation. Q Rev Biol 58(2):155−183.

Wieczorek A, Drewes R, Channing A (2000) Biogeography and evolutionary history of *Hyperolius* species: application of molecular phylogeny. J Biogeogr 27:1231–1243.

Wollenberg KC, Glaw F, Meyer A, Vences M (2007) Molecular phylogeny of Malagasy reed frogs, *Heterixalus*, and the relative performance of bioacoustics and color-patterns for resolving their systematics. Mol Phylogenet Evol 45:14–22.

Woolbright LL (1983) Sexual selection and size dimorphism in anuran Amphibia. Am Nat 121:110–119.

Yovanovich CAM, Koskela SM, Nevala N, Kondrashev SL, Kelber A, Donner K (2017) The dual rod system of amphibians supports colour discrimination at the absolute visual threshold. Phil Trans R Soc 372:20160066.

Zhang C, Sayyari E, Mirarab S (2017) ASTRAL-III: Increased Scalability and Impacts of Contracting Low Support Branches. Comparative Genomics. RECOMB-CG 2017. Lecture Notes in Computer Science, vol 10562, eds Meidanis J, Nakhleh L (Springer, Cham), pp 53−75.

